# Direct correlation between motile behavior and protein abundance in single cells

**DOI:** 10.1101/067918

**Authors:** Yann S Dufour, Sébastien Gillet, Nicholas W Frankel, Douglas B Weibel, Thierry Emonet

## Abstract

Understanding how stochastic molecular fluctuations affect cell behavior requires the quantification of both behavior and protein numbers in the same cells. Here, we combine automated microscopy with *in situ* hydrogel polymerization to measure single-cell protein expression after tracking swimming behavior. We characterized the distribution of non-genetic phenotypic diversity in *Escherichia coli* motility, which affects single-cell exploration. By expressing fluorescently tagged chemotaxis proteins (CheR and CheB) at different levels, we quantitatively mapped motile phenotype (tumble bias) to protein numbers using thousands of single-cell measurements. Our results disagreed with established models until we incorporated the role of CheB in receptor deamidation and the slow fluctuations in receptor methylation. Beyond refining models, our central finding is that changes in numbers of CheR and CheB affect the population mean tumble bias and its variance independently. Therefore, it is possible to adjust the degree of phenotypic diversity of a population by adjusting the global level of expression of CheR and CheB while keeping their ratio constant, which, as shown in previous studies, confers functional robustness to the system. Since genetic control of protein expression is heritable, our results suggest that non-genetic diversity in motile behavior is selectable, supporting earlier hypotheses that such diversity confers a selective advantage.

## Introduction

Cell behavior is controlled by biochemical signaling networks. Because network dynamics depend on the number of each protein involved, the dynamical and functional properties of signaling pathways—and ultimately cell survival—are sensitive to random fluctuations in protein numbers. Such fluctuations are expected because the molecular components of signaling pathways are typically present in small numbers [1]. Various mechanisms that cause molecular fluctuations have been identified, including stochastic protein expression or unequal partitioning of components between daughter cells [2, 3].

Selective pressure to maintain robust performance against inherent molecular fluctuations is likely to have played a significant role in the evolution of biological networks [4–7]. In fluctuating environments, however, non-genetic phenotypic variability can be an advantageous strategy for clonal populations raising the possibility that the distribution of different phenotypes within the population has a functional role beyond the mean phenotype [2,8–13].

Investigating how variability in protein number controls the distribution of cell performances requires experiments that can measure both behavior and protein numbers in individual cells. At the same time, accurate characterization of the distribution of protein numbers, phenotypes, and performances in a population requires high-throughput experiments to obtain sufficient statistics. Fluorescent reporters paired with video-microscopy or flow cytometry have enabled the detailed characterization of phenotypic variability and cellular response dynamics in many systems [1, 14–19]. Microfluidics have given substantial control over experimental conditions, allowing researchers to probe single-cell responses to perturbations over extended periods of time [20–26]. However, due to the relatively long exposure times required to measure fluorescence signal from a small number of molecules, current experimental techniques are fundamentally limited by the requirement that cells to be stationary, making it difficult to correlate fluorescent reporters directly with cell behaviors in the same cells.

We developed a high-throughput experimental method, FAST (Fluorescence Analysis with Single-cell Tracking), to correlate directly the individual behaviors of freely swimming cells and the numbers of proteins in the signaling pathway of each cell. FAST overcomes the conflicting requirements of tracking cell motile behavior over a large field of view with fluorescence imaging of stationary cells at high magnification. FAST uses *in situ* hydrogel polymerization to ‘freeze’ cells in place after the tracking period to allow automated single-cell fluorescence imaging. Therefore, the navigational performance of individual cells, such as exploration of an environment, was directly measured as a function of long-term motile behavior and intracellular protein numbers.

Here, we demonstrate FAST on the chemotaxis pathway of *Escherichia coli*, a model system for the study of cellular behavior. Motile *E. coli* cells explore their environment by alternating periods of relatively straight swimming (runs) with brief changes of direction (tumbles). Tumbles are caused by a reversal of the flagellar motor rotation from counter-clockwise to clockwise direction. The activity of the chemotaxis receptor cluster controls the frequency at which the cell tumbles by controlling the kinase activity of CheA, which phosphorylates the response regulator CheY. The probability to tumble — the tumble bias — increases when the phosphorylated form of CheY (CheY-P) binds the motor. The kinase activity decreases when the receptors bind attractant. As a result, the CheY-P concentration is decreased by the constitutive phosphatase CheZ and the tumble bias is reduced. If attractant concentrations remain steady, tumble bias returns to pre-stimulus level due to the activity of CheR and CheB. CheR methylates inactive receptors, which increases the activity of the kinase, whereas CheB demethylates active receptors, reducing kinase activity [27]. Theoretical studies predict that this resting tumble bias is an important determinant of the chemotactic performance of *E. coli* and that the fitness of a population of cells might depend not only on its mean tumble bias but also on the population variability in tumble bias [12, 13].

Early studies revealed a substantial amount of cell-to-cell variability in the motility behavior of clonal cells adapted to a uniform environment [28, 29]. However, the molecular origin and functional consequences of this variability remain unclear. Because CheR and CheB are engaged in a futile cycle of methylation and demethylation of the chemoreceptors, the balance between the actions of CheR and CheB is an important factor in determining the resting activity of the receptor cluster [27]. Consequently, variations in the numbers of CheR and CheB are expected to affect the tumble bias of single cells. How changes in the levels of expression of CheR and CheB affect the distribution of the tumble bias in a clonal population of cells is unknown. It also remains unclear if the mean and the variance of the tumble bias distribution can be controlled independently from each other. Here, we examine these questions experimentally by quantifying *in vivo* how variations in the numbers of CheR and CheB in single cells shape the distributions of swimming phenotypes and exploration capabilities *E. coli*.

## Results

### Cell-to-cell variability in motile behavior within a clonal population of *E. coli* cells

We recorded trajectories of freely swimming cells to determine the distribution of swimming phenotypes in an isogenic population. Following established protocols, *E. coli* RP437 cells were grown to mid-exponential phase in minimal medium then suspended in motility buffer in which the motile behavior remains constant for more than one hour and identical to the behavior observed in the growth medium (S1 Fig.) [30]. Therefore, the characterization of single-cell motile behavior was done without confounding effects from cell growth or protein turnover (Methods). Cells were imaged at low-density using phase contrast microscopy at 10X magnification in an isotropic liquid environment. Individual trajectories were reconstructed over a large field of view using a multiple-particle tracking algorithm [31] to enable the quantitative characterization of individual cell behavior (Fig. 1A, S2 Fig.). We constructed a probabilistic model using the instantaneous velocity, acceleration, and angular acceleration to describe the behavior of each cell along their trajectories and identify tumbling events (Fig. 1B, S3 Fig. AB). We then calculated the tumble bias (defined as the time spent tumbling over the total trajectory time) of each cell. The average cell behavior obtained from more than 6,000 cell trajectories was consistent with previously published, population-averaged results from *E. coli* RP437 cells. Specifically, the average tumble bias was 0.24 (Fig. 1C) with an average swimming speed of 33 μm/s (S3 Fig. D) [29, 32, 33].

**Fig. 1.**
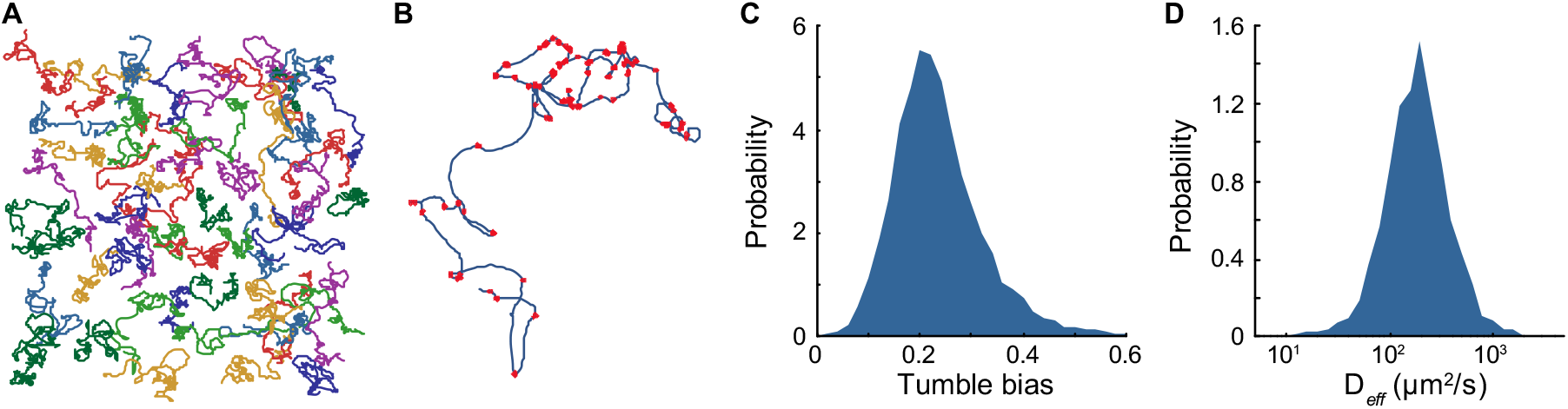
Measuring phenotypic diversity in motile behavior of *E. coli* RP437. (A) Representative trajectories of cells swimming in a liquid environment recorded using phase contrast microscopy at 10X magnification in a pseudo-2D environment. (B) Example of a 60 seconds single-cell trajectory where detected tumbles are marked with red dots. (C) Probability distribution of cell tumble biases in the isogenic population. (D) Probability distribution of individual cell diffusion coefficients (*D_eff_*) in the population. The distributions were calculated from about 6,000 individual trajectories combined from three independent experiments.

Single-cell trajectories revealed the variability in behavior within the isogenic population. The distribution of tumble biases was unimodal with mode at 0.2 and a standard deviation of 0.093 (Fig. 1C). These values are consistent with the previously reported distribution of single flagellar motor biases from tethered cells [29] and taking into account the effect of multiple flagella that increases the cell tumble bias relative to the clockwise bias of single motors [34, 35]. As expected, we observed very few cells with tumble biases outside the 0.1 to 0.4 range because the robust architecture of the chemotaxis pathway ensures that the population tumble bias is maintained within a functional range [7,36–40]. However, our experiment shows that there is still substantial phenotypic diversity among cells consistent with prior observations [28].

Phenotypic variability in tumble bias and swimming speed can result in large differences in how efficiently cells explore their environment. We calculated the diffusion coefficient of each cell by simultaneously fitting the mean square displacement and the velocity autocorrelation (S3 Fig. EFG) of the trajectories to the Green-Kubo relations [41, 42]. This analysis revealed that the effective diffusion coefficients of individual cells vary over more than one order of magnitude within the isogenic population in spite of the pathway robustness (Fig. 1D).

### Controlling motile behavior using fluorescently labeled CheR and CheB

The tumble bias of individual cells is determined in part by the numbers of the two receptor modification enzymes: CheR and CheB [12,29,38,43]. It has been previously observed that ectopic expression of CheR increases the cell tumble bias on average [29, 38]. Transcriptional and translational coupling in the expression of CheR and CheB provides robustness to the system against uncorrelated stochastic fluctuations in protein numbers to maintain good chemotactic performance [7, 44]. In addition, because of the slow exchange of the modification proteins between the cytoplasm and the cluster of receptors, the numbers of CheR and CheB affect signaling noise, which results in slow fluctuations of the cell tumble bias [29, 45, 46]. To understand the consequences of variations in the numbers of CheR and CheB on the swimming phenotype of *E. coli*, we sought to make a detailed quantitative map of motile behavior as a function of protein numbers.

We genetically constructed the fluorescent protein fusions CheB-mYFP and mCherry-CheR, which have been previously shown to be functional proteins [47]. To explore a large dynamic range of absolute protein expression and CheR to CheB ratio, the gene constructs were placed under the control of two independent inducible promoters on the chromosome in a strain lacking both the native *cheR* and *cheB* genes (Fig. 2A). The separate transcriptional regulation of the two genes allowed for the control of both the absolute numbers and the ratio of CheB-mYFP and mCherry-CheR (Fig. 2BC, S4 Fig.). The fluorescence intensities per molecule of CheB-mYFP and mCherry-CheR was calibrated using quantitative immunoblotting and single-cell fluorescence microscopy [48], following a previously described method [49]. The fluorescence intensity per fluorescent protein was calculated using a Bayesian regression analysis (Methods and S5 Fig.).

**Fig. 2.**
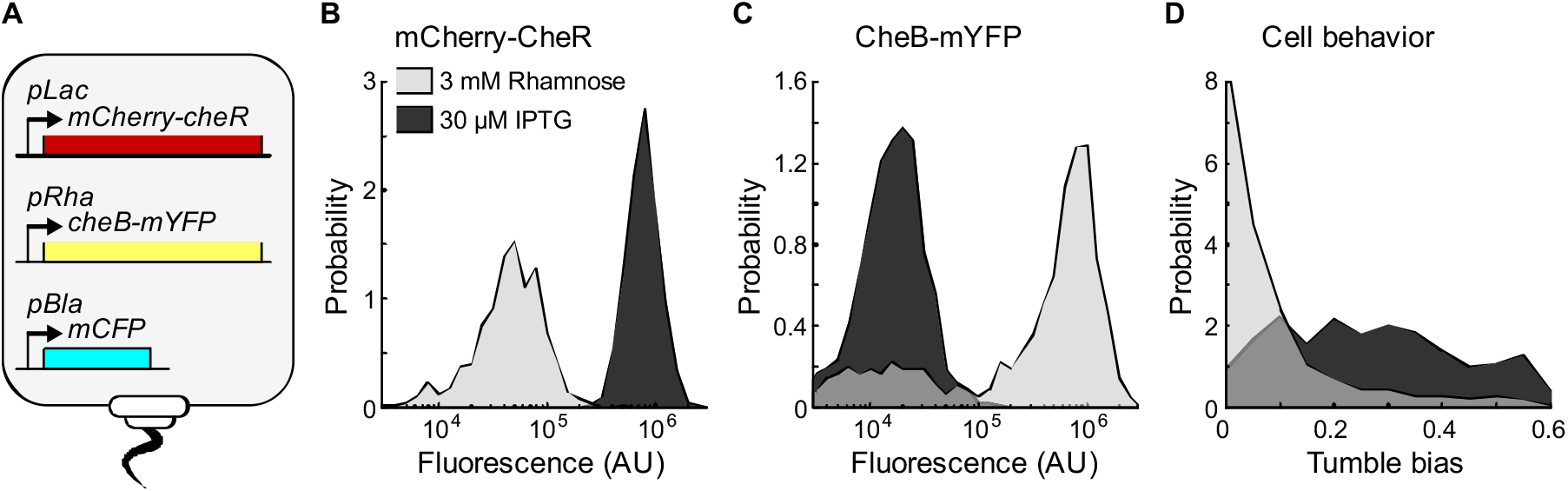
Controlling motile behavior using fluorescently labeled CheR and CheB. (A) Diagram representing a mutant strain derived from *E. coli* RP437 *DcheRcheB* expressing fluorescent protein fusions from inducible promoters recombined in the cell genome. (B) Distributions of red fluorescence intensity from single-cell epi-fluorescence microscopy from the differential inductions of mCherry-CheR with rhamnose (in light) or IPTG (in dark). (C) Distributions of yellow fluorescence intensity from single-cell epi-fluorescence microscopy from the differential inductions of CheB-mYFP with rhamnose (in light) or IPTG (in dark). (D) Distributions of single-cell motile behavior resulting from high expression of mCherry-CheR relative to CheB-mYFP (in dark) or high expression of CheB-mYFP relative to mCherry-CheR (in light).

**Fig. 3.**
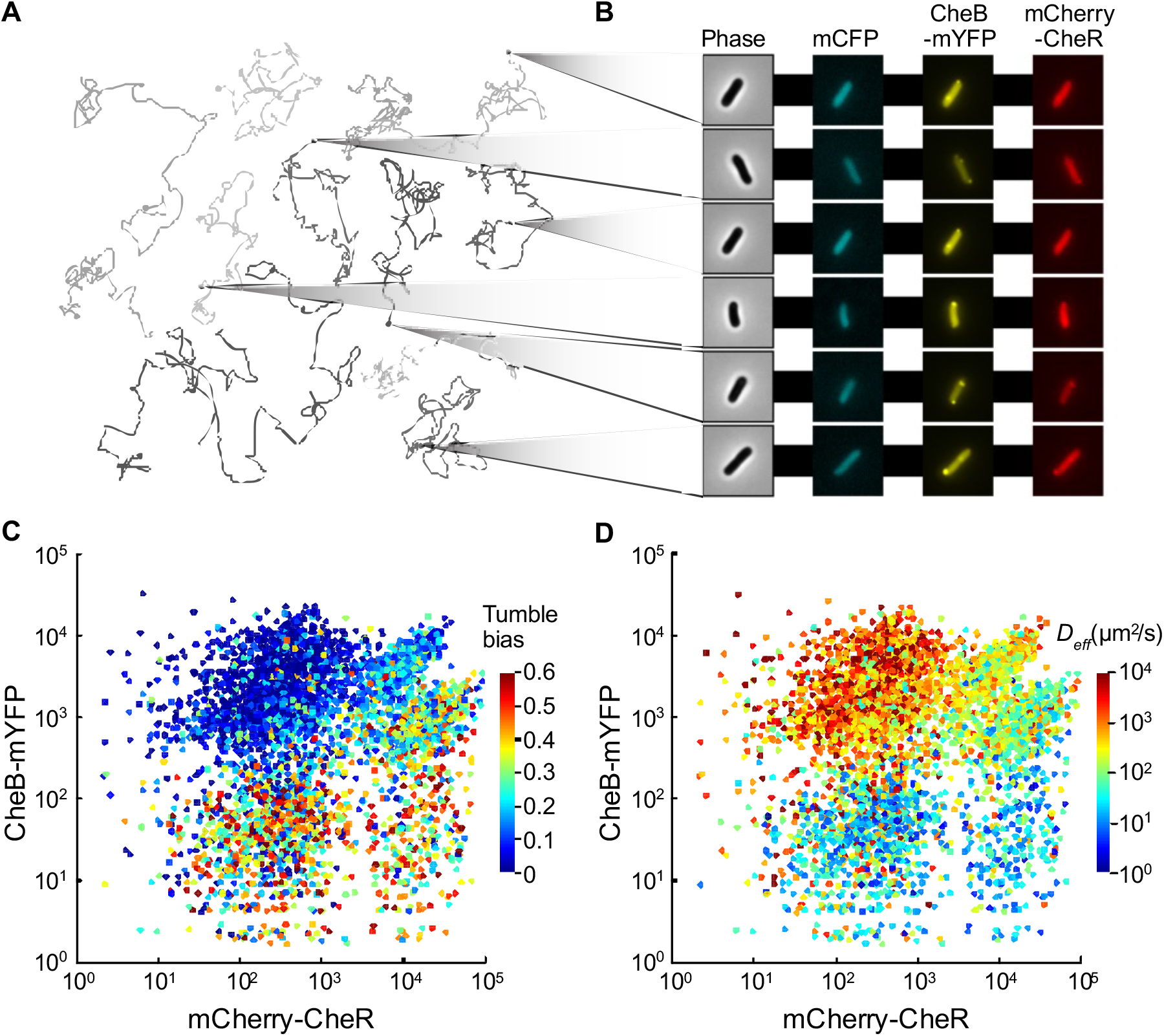
Mapping swimming phenotype to mCherry-CheR and CheB-mYFP numbers. (A) Representative trajectories of cells swimming in the non-polymerized hydrogel recorded using phase contrast microscopy at 10X magnification. (B) Examples of cells imaged using automated epi-fluorescence microscopy at 100X in three channels after cells were trapped in the polymerized hydrogel. (C) More than 4,000 cell trajectories were matched to fluorescence images to map the cell tumble bias of each cell as a function of mCherry-CheR and CheB-mYFP numbers. (D) The effective diffusion coefficients (*D_eff_* of individual cells as a function of mCherry-CheR and CheB-mYFP numbers. Multiple experiments using cells treated with different concentrations of inducers were combined.

We used the mutant strains to create different populations with widely different distributions of tumble bias (Fig. 2D, S6 Fig. and S7 Fig.). Consistent with previous work that measured average population behavior, the tumble bias distribution of cells increased with higher expression of mCherry-CheR and decreased with higher expression of CheB-mYFP, which has antagonistic activity to CheR [29,38,43,50]. The observed variability in protein expression and motile behavior within the population for each induction level is substantial despite the fact that each gene is expressed from a single chromosomal locus (Fig. 2D, S6 Fig. and S7 Fig.). Although CheR and CheB expression in the mutant strains is different from the wild type regulation, the results illustrate in general how phenotypic diversity can be altered through changes in the activity of single promoters.

### Direct mapping of swimming phenotype to proteins numbers

The FAST protocol relies on the immobilization of cells in a hydrogel, which is polymerized *in situ*, after tracking motile cells. We first mixed cells washed with motility buffer with a hydrogel precursor solution that contained polyethylene glycol diacrylate (PEGDA) and a photoinitiator (LAP) [51]. The presence of PEGDA and LAP did not affect motile behavior beyond a small reduction in swimming speed (S1 Fig.). Swimming was recorded at 10X as described above for 5 minutes (Fig. 3A). A flash of violet light was delivered through the microscope objective to activate the photo-initiator and trigger rapid hydrogel polymerization over the entire field of view. Cells were immobilized within 1 second and their coordinates were registered by processing the real-time image from the camera. After switching the microscope configuration for fluorescence imaging at 100X magnification, each cell was automatically located by a motorized stage and imaged in phase contrast and three fluorescence channels (Fig. 3B). An average of 200 cells was imaged in less than 40 minutes for each experimental trial. The fluorescence signal from each cell did not change significantly as a function of time during the single-cell imaging phase indicating that the fluorescent protein fusions are stable when the cells are trapped in the hydrogel (S8 Fig.). Therefore, the measured single-cell fluorescence corresponds to the number of proteins present while the cells were swimming. Finally, cells trajectories were matched to fluorescence images in order to map motile behavior to proteins numbers in single cells.

We combined measurements from several experiments in which different combinations of inducer concentrations were used with the two fluorescent strains to produce a wide range of phenotypes, with cells expressing the labeled proteins over three orders of magnitude and tumble bias ranging from 0 to 0.6. We mapped tumble bias to the number of mCherry-CheR and CheB-mYFP for more than 5,000 individual cells (Fig. 3C). Tumble bias increases with higher numbers of mCherry-CheR and decreases with lower numbers of CheB-mYFP, consistent with population measurements. We verified that the phenotypes showed no correlation with the constitutively expressed mCFP protein that was not fused to any of the native *E. coli* proteins (S9 Fig.). The uneven sampling of CheR and CheB (Fig. 3C) is due to bistability of the activity of the rhamnose promoter that was used to control protein expression. We also mapped the individual diffusion coefficients to mCherry-CheR and CheB-mYFP numbers (Fig. 3D). Because of the large range of protein expression in this experiment, the individual cell diffusion coefficients span four orders of magnitude. This variability is larger than what was observed in the wild-type population (Fig. 1C), indicating that the natural cell-to-cell variability in CheR and CheB numbers is constrained to a range smaller than the artificially induced conditions. As expected, the relationship between the cell diffusion coefficients and protein numbers (Fig. 3D) mirrors the relationship obtained from characterizing the cell tumble biases (Fig. 3C). Because the value of diffusion coefficient calculated from a trajectory does not rely on tumble detection, this analysis provides additional support for the observed effect of variations in protein numbers on behavior.

### The mean and the variance of the tumble bias are affected in different ways by changes in mCherry-CheR and CheB-mYFP numbers

Together, the single-cell data provide a direct mapping of protein numbers to swimming phenotype with unprecedented detail over a large range of protein numbers. We first examined how CheB-mYFP and mCherry-CheR control the mean tumble bias by performing a local linear regression fit of the single-cell tumble bias (Fig. 4A). The relationship between mean tumble bias and the logarithm of the CheB-mYFP and mCherry-CheR numbers has the characteristic feature of diagonal contour lines of increasing tumble bias, qualitatively indicating that tumble bias depends more on the ratio than the absolute number of these proteins. This trend was expected since CheR and CheB have antagonistic effects on kinase activity by participating in a futile cycle of methylation and demethylation of the chemoreceptors. However, the tumble bias appears to be more sensitive to changes in CheB-mYFP numbers than mCherry-CheR numbers. For example, changing the tumble bias by 0.1 requires approximately a 10-fold change in CheB-mYFP (Fig. 4B) but a 40-fold change in mCherry-CheR (Fig. 4C). We analyze this unexpected asymmetry in more detail in the next section.

**Fig. 4.**
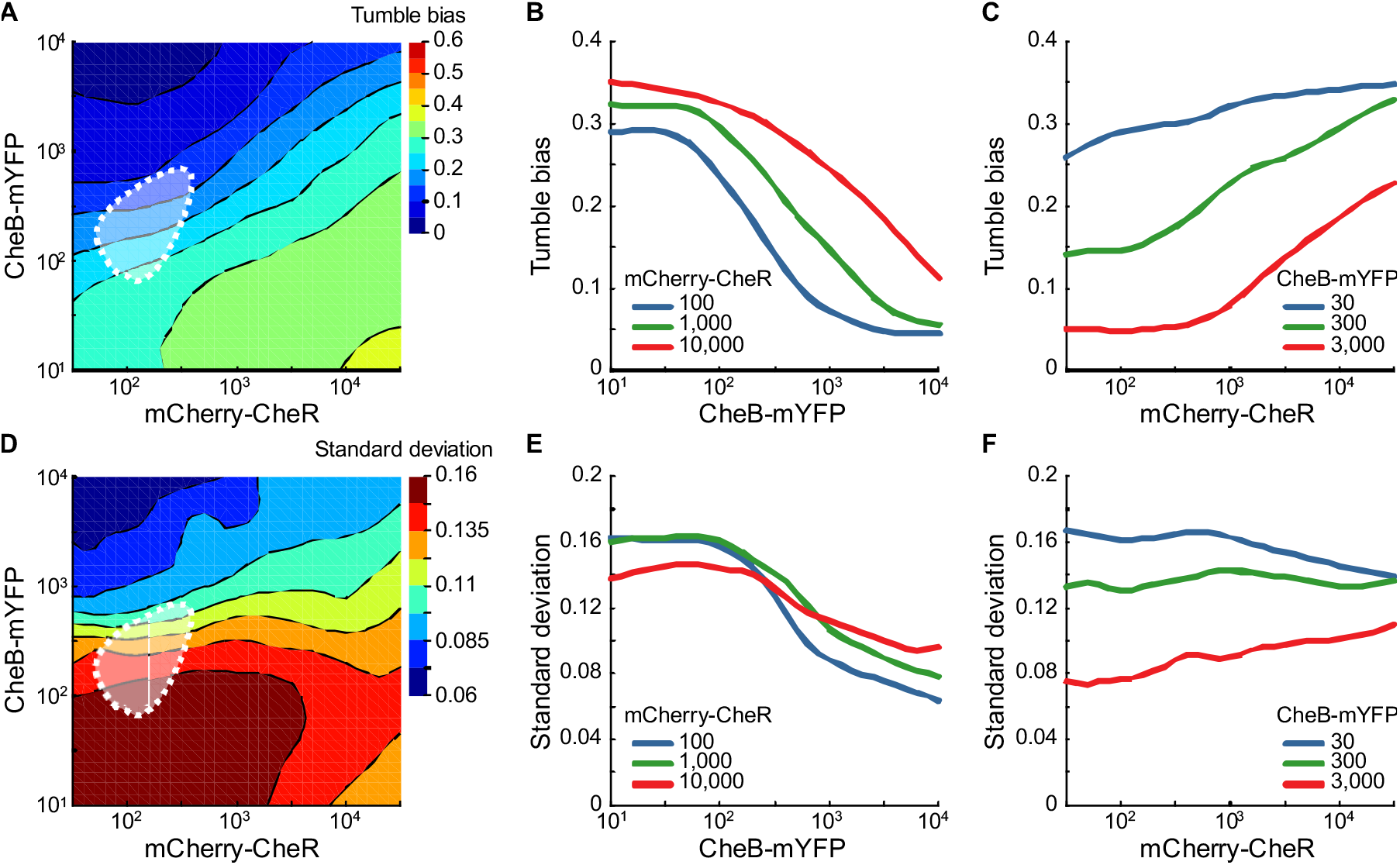
Analysis of phenotypic variability. (A) Contour plot of the local linear regression of the tumble bias as a function of mCherry-CheR and CheB-mYFP numbers. The color scale is the same as Fig. 3C. (BC) Cross-sections of the local linear regression of the tumble bias showing the relative sensitivity of the mean tumble to changes in protein numbers. (D) Contour plot of the local linear regression of the residual tumble bias variance after subtracting the change in tumble bias explained by mCherry-CheR and CheB-mYFP numbers. (EF) Cross-sections of the local regression of the variance showing the relationship between residual phenotypic diversity and CheB-mYFP or mCherry-CheR numbers. The expected cell-to-cell variation in CheR and CheB numbers in the wild-type population is indicated by the white dotted contour (95% of the probability density function) [12, 44, 52]. The local linear regressions were done using a bandwidth of 20% of the data points.

With a quantitative relationship between tumble bias and the numbers of CheB-mYFP and mCherry-CheR, we asked how much of the variability in tumble bias measured in the wild-type population (Fig. 1E) could be attributed to the natural cell-to-cell variability in the numbers of CheR and CheB. In wild type cells, *cheR* and *cheB* are transcriptionally and translationally coupled, hence fluctuations in protein numbers have correlated lognormal distributions with extrinsic noise (*η_ext_*) and a smaller intrinsic noise (*η_int_*). This genetic organization prevents large variations in tumble bias as a result of fluctuations in protein numbers [7, 44]. Using previously reported estimates of the noise intensity in the expression of the chemotactic proteins, we generated the expected distributions for CheR and CheB numbers across individual cells in the wild type population (Fig. 4A). Then, we used the quantitative relationship between tumble bias and protein numbers to estimate the contribution of the natural cell-to-cell variability in CheR and CheB numbers to the observed cell-to-cell variability in tumble bias in the wild-type population. Using *η_ext_* = 0.26 and *η_int_* = 0.125 [12, 44, 52], CheR and CheB fluctuations explain 11% of the variance in tumble bias. Therefore, additional factors must contribute to a large portion of the wild type variability in tumble bias. Likely candidates are variations in the numbers of CheA, CheY, CheZ, the number of flagellar motors (cell-to-cell variations), or slow stochastic fluctuations in the methylation levels of the receptors (single-cell variations) [12, 45].

Further examination of the single-cell data indicates that there is large residual variability around the mean tumble bias for any given level of mCherry-CheR and CheB-mYFP. Calculating the standard deviation of the residual tumble bias as a function of mCherry-CheR and CheB-mYFP (Fig. 4D) reveals that the residual variability depends strongly on CheB-mYFP (Fig. 4E) and weakly on mCherry-CheR (Fig. 4F). Remarkably, the dependency of the residual variability on mCherry-CheR and CheB-mYFP numbers is not aligned with the dependency of the mean tumble bias (the contours in Fig. 4A and 4D are not aligned in the same direction). This observation suggests that the mean and the variance of the tumble bias distribution can be adjusted independently from each other by controlling chemotaxis protein expressions. Importantly, focusing on the data along the diagonal in Fig. 4AD reveals that the amount of phenotypic diversity in the population can be adjusted by changing the global level of expression of CheR and CheB while simultaneously maintaining their ratio nearly constant, therefore maintaining the robustness of chemotaxis pathway conferred by the co-expression of CheR and CheB from one operon [44, 53]. These findings support the hypothesis that phenotypic diversity in *E. coli* chemotaxis is a selectable trait [12, 54].

### Modeling the mean and the variance of the tumble bias as a function of CheR and CheB numbers

Current standard models of the bacterial chemotaxis system do not explain two aspects of our experimental results [12, 54]. First, the observed mean tumble bias is not equally sensitive to changes in CheB-mYFP and mCherry-CheR numbers. A fit of the tumble bias to the logarithm of the protein numbers gives the relationship: 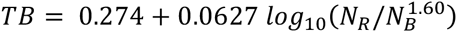, where *N_R_* and *N_B_* are the numbers of mCherry-CheR and CheB-mYFP proteins in a single cell (the 95% confidence intervals for the parameters are, in order, [0.258, 0.291], [0.0586, 0.0668], and [1.49, 1. 72]). We investigated whether the secondary feedback loop in chemotaxis system (created by the phosphorylation of CheB by CheA) could increase the sensitivity of the tumble bias to CheB numbers. We found that the tumble bias would still be determined by a simple ratio of CheR to CheB numbers consistent with previous models [12,43,54,55] (S10 Fig.).

Second, the variance in tumble bias does not depend on mCherry-CheR and CheB-mYFP numbers the same way the mean tumble bias does (Fig. 4C and 4F). This observation was unexpected because it is not explained by models that assume that cell-to-cell variability in tumble bias results solely from cell-to-cell variability in chemotaxis protein numbers. Published models predict that the mean and the residual variance of the tumble bias have parallel dependencies on variations in CheR and CheB numbers [12, 56] (S10 Fig. illustrates the theoretical predictions from these models, wherein the contours of mean and variance in tumble bias are aligned in the same direction).

To explain our experimental findings, we propose a model that takes into account the dual role that CheB plays in both demethylating and deamidating the receptor proteins [57] and the slow fluctuations of the methylation level of the receptor cluster [45, 46]. When the most abundant receptors, Tar and Tsr, are synthesized, they are translated with two glutamines (Q) instead of glutamates (E) at the methylation sites in a QEQE configuration. Glutamate residues can be reversibly methylated and demethylated to adjust the basal activity, or free energy, of the receptors. Non-mature receptors have a semi-active conformation in the absence of stimuli and cause higher-than-expected tumble bias in a *cheRcheB* mutant [58] because glutamine acts similarly to a methylated glutamate but with half the change in receptor free energy [43, 59, 60]. CheB irreversibly deamidates the glutamines to glutamates so that the residues can then be used for adaptation [57]. Therefore, cells need to synthesize and deamidate a full set of receptors during each cell division to ensure that all modification sites are available for reversible methylation.

We hypothesized that high tumble bias arising from low levels of CheB-mYFP is caused by the incomplete deamidation of the receptors. We introduce a deamidation rate equation to take into account the maturation of receptors by CheB:

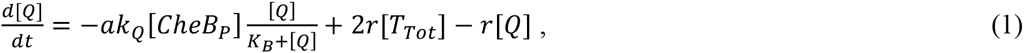

where *a* is the activity of the receptor cluster, [*CheB_p_*], [*T_Tot_*], and [Q] are the concentrations of phosphorylated CheB, total receptors, and glutamine residues, *k_Q_* is the deamidation rate, *K_B_* is the Michaelis-Menten constant characterizing the CheB-receptor binding, and *r* is the cell growth rate. The first term in equation (1) corresponds to the rate of CheB-dependent deamidation, and the last two terms correspond to generation and dilution of glutamine residues within a cell as new receptors proteins are synthesized and cell divides. We modified the (de)methylation rate reaction [12] accordingly to take into account the presence of glutamine residues that cannot be (de)methylated and the dilution of methylated receptor by cell growth:

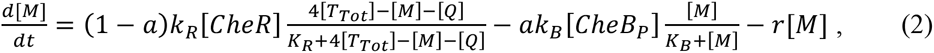

where [*CheR*] and [M] are the concentrations of CheR and methylated glutamate residues, *k_R_* and *k_B_* are the methylation and demethylation rates, *K_R_* is the Michaelis-Menten constant characterizing the CheR-receptor binding. The first term in equation (2) corresponds to the rate of CheR-dependent methylation of the available glutamate residues, the second term corresponds to the CheB-dependent demethylation of the methylated glutamate residues, and the last term corresponds to the dilution of the methylated residues from cell growth. The remaining equations describing the dynamics of phospho-relay reactions remain unchanged [12] because the kinetics rates remain much faster than the cell growth rate even when CheR and CheB numbers are low (see Methods). Finally, the activity of the receptor in the absence of chemical stimuli as a function of glutamine and methylated glutamate residues is described by:

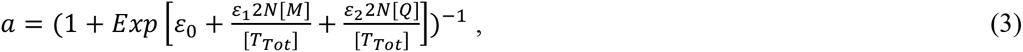

where *ε_0_ ε_1_* and *ε_2_* are free energy constants, *N* is the size of the MWC complexes [54]. In the absence of chemotactic signals, the system can be solved at equilibrium to determine the steady-state tumble bias. To simulate cell-to-cell variability in protein expression we sampled protein numbers for each cell from lognormal distributions with intrinsic and extrinsic noise generators [12].

Introducing the effects from the synthesis of non-mature receptors and their deamidation by CheB in the model was sufficient to reproduce the higher sensitivity of the cell tumble bias to CheB than CheR (Fig. 5A and S10 Fig. C, the orientation of the contours produced by our model matches the data in Fig. 4A). The model predicts that when the number of CheB molecules becomes limiting, glutamine residues accumulate in the receptor cluster during cell growth resulting in an increase of the average tumble bias. We verified this hypothesis by following the population tumble bias immediately after transfer from growth medium to chemotaxis buffer. When the number of CheB is lower than the wild-type population mean of 240 molecules per cell [52], the population average tumble bias starts high and slowly decreases over the course of an hour (S11 Fig.). From this experiment, we estimated that the CheB-dependent deamidation rate is half of the demethylation rate.

**Fig. 5.**
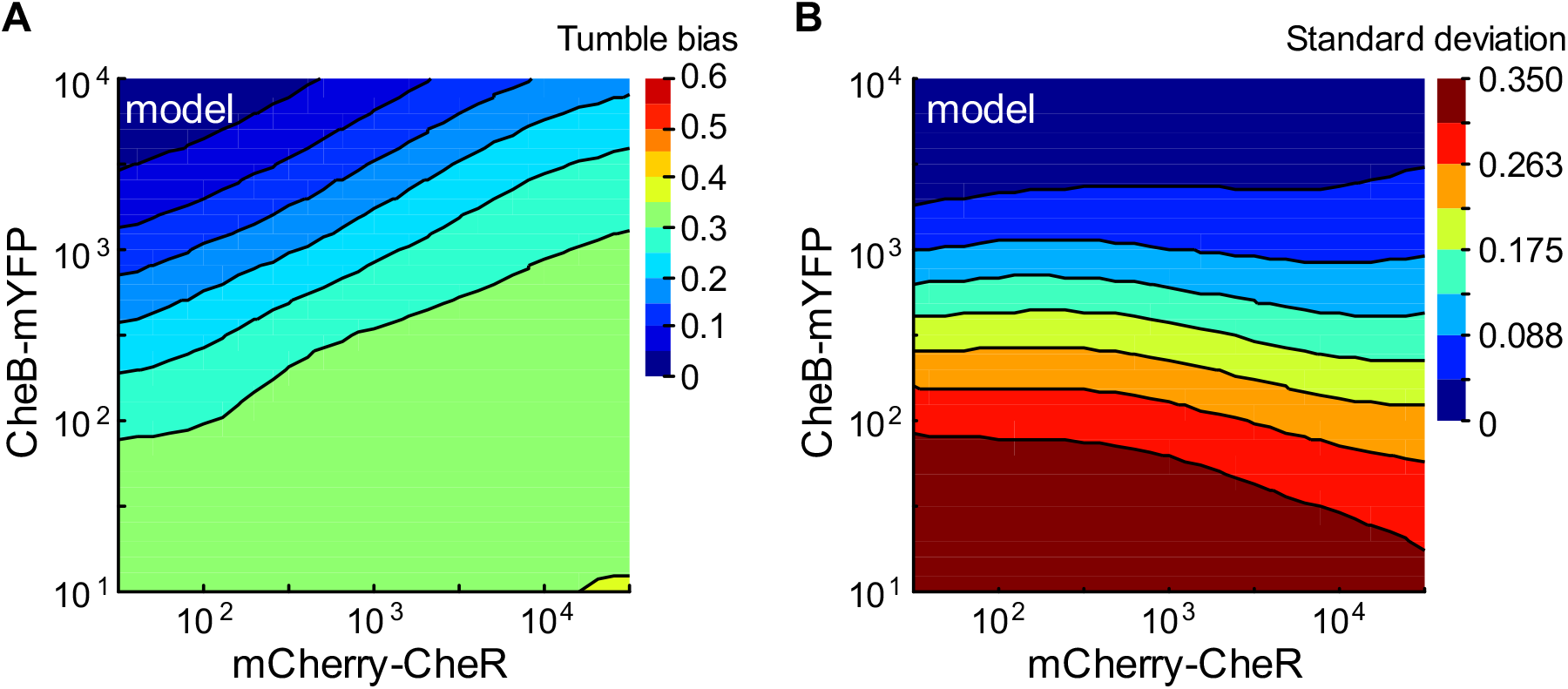
Modeling the mean and variance of the tumble bias as a function of CheR and CheB numbers. (A) Contour plot of the local linear regression of the predicted tumble bias as a function of CheR and CheB numbers. (B) Contour plot of the residual tumble bias standard deviation after subtracting the change in the predicted tumble bias explained by CheR and CheB numbers. The tumble bias and adaptation time were calculated according to the modified chemotaxis model for 8405 cells covering the full range of CheR and CheB numbers. The local linear regressions were done using a bandwidth of 20% of the data points.

To explain the relationship of the residual variance of the tumble bias as a function of CheR and CheB, we took into account the slow fluctuations of the methylation level of the receptor cluster within the timescale of our experiments. When the numbers of CheR and CheB are small compared to the number of receptors, the average methylation level becomes hyper-sensitive to the ratio of the two modification proteins [45, 46, 61]. As a result, the tumble bias of one cell can fluctuate significantly over time scales similar to the duration of the tracks that we used to quantify tumble bias in our experiment [29, 46]. When the spontaneous fluctuations of the receptor activity are taken into account, the residual variance in tumble bias becomes more dependent on CheB rather than the mean tumble bias (Fig. 5B, the orientation of the contours produced by our model align with changes in CheB numbers similar to what is observed in the data in Fig. 4D). Overall, the dependency of the observed residual variance supports the hypothesis that behavioral variability in a clonal population is a result of both signaling noise caused by the receptor adaptation dynamics and cell-to-cell variability in protein numbers.

## Discussion

Understanding the functional role of variability in clonal populations of cells will require understanding how molecular variations map onto phenotypic variations, which in turn translate into performance differentials between individual cells. While molecules, phenotype, and performance of individual cells can all be measured separately [17,28,29,33,46,62,63], making all these measurements in the same cells has not been possible due to the large differences in length scales and time scales involved. By combining fast *in situ* hydrogel polymerization with automated fluorescence microscopy, we were able to bridge scales and directly correlate for the first time individual motile behaviors of freely swimming cells to intracellular protein numbers.

We mapped single-cell tumble bias and exploratory capability as a function of the numbers of the two adaptation proteins of the chemotaxis pathway, CheR and CheB, with unprecedented details. We found that CheR and CheB numbers affect both the mean and the variance of the tumble bias but in different ways. Therefore, the shape of the phenotypic distribution in an isogenic population could be adjusted through genetically encoded factors such as the levels of protein expression. This suggests that the variability in tumble bias can evolve in an isogeneic population while the mean tumble bias remains constant and *vice versa* solely through mutations that change the relative expression levels of CheR and CheB (and possibly other chemotaxis proteins such as CheY and CheZ). These experimental observations support previous theoretical predictions that the degree of phenotypic diversity in swimming behavior could be a selectable trait in *E. coli* [12, 13]. Previous studies demonstrated that translational and transcriptional coupling of CheR and CheB confer robustness to the chemotactic system [44, 53] and that even when phenotypic diversity is advantageous it is important to maintain specific ratios in the numbers of proteins [12]. Our results show that a clonal *E. coli* population can adjust phenotypic diversity by adjusting the total expression CheR and CheB without disrupting their coupling. We also found that single-cell behavioral variability caused by the dynamics of receptor methylation, as previously described [45, 46], contributes significantly to the observed population phenotypic diversity in addition to cell-to-cell variability in protein expression. The contributions of additional molecular and morphological factors, such as the number of flagella, cell shape, or the location of the receptor clusters, to individual cell motile behavior remain to be characterized. By enabling the quantitative measurement of multivariate distributions, FAST will facilitate the characterization of phenotypic variability as a function of protein numbers, signaling pathway architecture, or other cell components.

Taking advantage of the large field of view and high-resolution offered by modern scientific cameras, we were able to track and quantify the tumble of thousands of wild type *E. coli* cells.

We found that tumble bias and diffusion coefficient were widely distributed. This variability is expected to have a significant impact on the spatial organization and fitness of cells when competing for resources [12,64–66]. Few cells had tumble biases above 0.4, consistent with predictions that high tumble biases perform poorly [13].

To explain the unpredicted finding that tumble bias is more sensitive to CheB than CheR, we propose that the deamidation of the newly synthesized receptor proteins becomes incomplete when the number of CheB falls below approximately one hundred molecules, which is within reason since the mean expression level is ~240 [52]. With incomplete deamidation, the basal activity of the receptor is increased, causing elevated tumble bias not explained by previous models. From the analysis of our model, we found that the activation of CheB through phosphorylation by CheA was not sufficient to explain our experimental observations because this feedback does not introduce an asymmetry in the relationship between the mean and the variance of the tumble bias as a function of CheR and CheB numbers.

The biological significance of the CheB-dependent maturation of the dominant receptor proteins via the deamidation of specific Q residues is still not understood. However, our results suggest that wild-type cells express on average just enough CheB to keep up with the synthesis and maturation of new receptors during growth. One possibility is that the QEQE configuration may place the (de)methylation dynamics of the receptor cluster closer to equilibrium, saving a significant amount of time and energy to methylate new receptor proteins since they outnumber CheR and CheB by about two orders of magnitude.

Another possibility is that synthesizing receptor proteins with a QEQE configuration is a bet-hedging strategy. Because of cell-to-cell variability in the expression levels of CheR and CheB some cells will express few (de)methylation enzymes. Previous work has shown that when CheR and CheB are limiting the tumble bias become hyper-sensitive to the ratio of the numbers of CheR and CheB [45, 46, 61]. Upon expression of the chemotaxis pathway cells should initially have higher tumble bias and therefore stay close to their sisters due to the higher activity of the QEQE configuration. However, as the chemotaxis receptors become fully deamidated, individual cells will start leaving the colony and explore their surroundings. Cell-to-cell variability in the expression of CheR and CheB may result in a slow trickling of explorers from the colony. This slow transition from tumbler to explorer may be a bet-hedging strategy when turning the chemotaxis pathway on.

An important aspect of signal transduction is that changes in behavior affect how cells interact with environmental signals. This is especially true for navigation where behavior feeds back onto the statistics of input signals and *vice versa* [13]. This important feedback loop is lost when cell are attached on surfaces or immobilized with optical tweezers [35, 40, 59]. FAST alleviates these constraints, by allowing behavioral tracking and fluorescence imaging of freely swimming cells. The combination of FAST with the use of nano-fabricated landscapes to create chemical gradients should facilitate the investigation of the molecular, cellular, and population level mechanisms that underlie the emergent behaviors of cells in complex environments.

## Methods

### Strains and growth conditions

*E. coli* RP437 was used as the wild-type strain for chemotaxis and as the parental strain for all the mutants generated in this study. Cells were cultured in M9 minimal medium supplemented with 10g/L glycerol, 1g/L tryptone, 2 mM magnesium sulfate, 0.1 mM calcium chloride, 10 mg/L thiamine hydrochloride, and 50 mg/L streptomycin. Isopropyl β-D-1-thiogalactopyranoside (IPTG) and rhamnose were added to the growth medium when indicated to induce protein expression. Cells were grown in aerobic conditions at 30°C in an Erlenmeyer flask on an orbital shaker at 200 rpm for aeration. Starting from single colonies isolated on agar plates, cells were grown to saturation overnight in broth cultures and sub-cultured using 1:100 dilution ratio in fresh medium and grown to an optical density at 600 nm (OD_600_) of 0.25. Under these growth conditions, virtually all cells were highly motile.

### Time-lapse microscopy and cell tracking

Before performing the behavioral experiments, cells were washed twice at room temperature with motility buffer (M9 salts supplemented with 0.1 mM ethylenediaminetetraacetic acid (EDTA), 0.01 mM L-methionine, 10 mM sodium lactate, and 0.05% weight/volume polyvinylpyrrolidone (M.W. ~40,000 Da)) by centrifuging cells at 2,000 g for 5 minutes and diluted to a low cell density (OD_600_ ~ 0.01). The buffer exchange and centrifugation did not appear to affect the cell behavior when compared to cells sampled from the growth medium (S1 Fig.). To record motile behavior, 5 μL of cells in motility buffer was sealed between a glass microscope slide and a 22 mm^2^ #1.5 coverslip using VALAP (equal amount of petrolatum, lanolin, and paraffin wax). Cells were free to swim in a pseudo two-dimensional environment ~10 μm deep. Cell motion was recorded at 10 frames per second with a digital scientific CMOS camera (Hamamatsu ORCA-Flash4.0 V2, 2x2 pixel binning, 50 ms exposure, rolling shutter, full frame) mounted on an inverted microscope (Nikon Eclipse TI-U) with a 10X phase contrast objective (Nikon CFI Plan Fluor, N.A. 0.30, W.D 16.0mm) and LED white light diascopic illumination (Thorburn Illumination Systems). The field of view was ~1.3 mm square containing on average 200 cells.

To reconstruct the cell trajectories each image sequence was processed using custom MATLAB (Mathworks) code. First, the mean pixel intensities of the frames over the entire image sequence was calculated to obtain an image of the background and subtracted from each image. The subpixel resolution coordinates of each cell in each frame were detected using a previously described method using radial symmetry [67] with an intensity detection threshold set to 6 standard deviations over the background. Coordinates were linked to obtain cell trajectories using a previously described self-adaptive particle tracking method, u-track 2.1 [31], with the linear motion model linkage cost matrices, an expected particle velocity of 30 μm/s, and otherwise default parameters.

### Cell trajectory analysis

The cell velocity at each time point was calculated according to 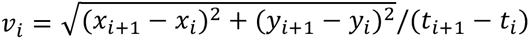. The acceleration was calculated according to 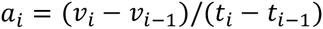. The angular acceleration was calculated according to *α_i_* = ((*θ_i_* – *θ_i−1_*) – (*θ_i−1_* – *θ_i−2_*))/(*t_i_*−*t_i−1_*)^2^. The velocity auto-correlation and mean square displacement of each trajectory were analyzed to extract the average mean run time and diffusion coefficient of each cell. The velocity autocorrelation, *C_v_*, and the mean square displacement, MSD, were calculated according to: 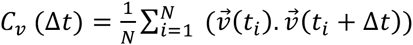 and 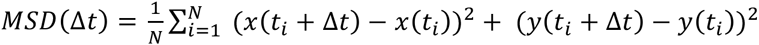, where *t_i_* represents the relative time for each frame of the image sequence, and Δ*t* the time interval between time points. The data was fitted using a non-linear least-square method with the functions: 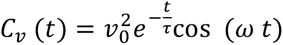, and 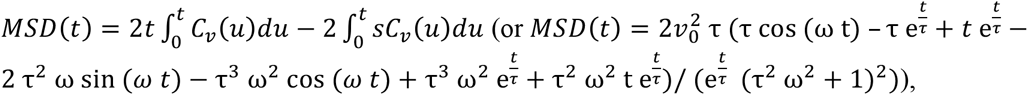, where *t* is time, *v*_0_ is the average cell speed, *τ* is the time scale of the cell directional persistence (a function of the cell tumble bias, mean tumble angle, and rotational diffusion), and *ω* is the angular frequency of the circular trajectory resulting from the interaction when cells swim near the glass surface (S12 Fig.). The mean square displacement function was calculated by taking the integral of the velocity autocorrelation function in two dimensions according to the Green-Kubo relations [41, 42]. The effective diffusion coefficient, *D*, was calculated according to 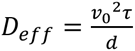 to remove the effect of the glass–surface interaction, where *d* is the number of dimensions (two in our experiment). The effective diffusion coefficient is in good agreement with a previously derived approximation of the diffusion coefficient derived for swimming *E. coli* cells as a function of the mean run time between tumbles [68] defined as: 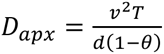, where *T* is the average run time between tumbles calculated for each cell using our tumble detection analysis, and *θ* is the mean cosine of the tumble angles (θ = 0.18 in our dataset, S3 Fig. H).

The posterior probabilities for a cell to be swimming (S), tumbling (T), and an intermediate state recovering from tumbling (I) given the instantaneous velocity (*v*), acceleration (*a*), and angular acceleration *(α)* (P(S|*v,a,α*), P(T|*v,a,α*), and P(I|*v,a,α*)) were constructed from a reference dataset containing more than 6,000 trajectories from wild-type cells. The parameters of each distribution were estimated by fitting a mixture of three tri-variate Gaussian distributions to the pooled distributions of instantaneous velocity (*v*), accelerations (*a*), and angular acceleration (*α*) (S3 Fig. AB). Therefore, each behavioral state is represented by a tri-variate Gaussian distribution. The mixture model was fitted to the reference dataset using an iterative approach. First, the swimming speed of each cell was normalized by their average speed when in the swimming state (the first iteration was initialized using the 95 ^th^ percentile of their instantaneous speeds). The relative acceleration and the angular acceleration between consecutive velocity vectors were computed. Then, all the relative speed, acceleration, and angular acceleration, were fitted with the mixture model. Each time point of the cell trajectories was assigned to the state with the largest posterior probability. The normalization, fitting, and state assignment were done iteratively until changes in state assignment between consecutive iterations converged below a tolerance of 1% (10 iterations on average). The resulting posterior probabilities were used to analyze the trajectories of all the cells in this work. Based on our validation of the tumble detection on simulated trajectories (see next section), we discarded all trajectories that were shorter than 10 seconds because short trajectories resulted in inaccurate tumble bias calculations (the code is available at https://github.com/dufourya/SwimTracker).

### Simulated cell trajectories

To validate the tumble detection model we simulated the swimming trajectories of cells with defined phenotypes in the absence of signal gradients. Simulations were run following a previously described method [34], with a constant swimming speed of 20 μm/s and rotational diffusion of 0.062 rad^2^/s [33]. Cells are stationary during tumbles and their orientations are uniformly randomized. The simulated cell tumble bias was changed by varying the internal CheY-P, *Y_p_*, concentration, which controls the transitions rates k_+_ and k___ to clockwise and counter clockwise of the flagellar motor according to: 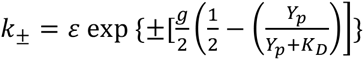, with *ε* = 1.3 s^−1^, *g* = 40, and *K_D_* = 3.06 μM, according to previously published experimental data [40]. The simulated environment was bounded in the z-dimension by reflecting boundaries 10 μm apart. Cell positions were sampled every 100 ms and projected in two dimensions to reproduce the experimental conditions. The accuracy and precision of the trajectory analysis was determined using 1,000 simulated trajectories for each tumble bias (S13 Fig.). The simulations showed that the tumble detection and tumble bias calculations were accurate for trajectories as short as 10 seconds.

### Construction of plasmids and mutant strains

Plasmids and mutant strains were constructed following standard cloning protocols (see S1 Table, S2 Table and S3 Table for the lists of plasmids, strains, and oligonucleotides). The deletion of *cheR* and *cheB* in *E. coli* RP437 was done using the λ Red disruption system [69]. Approximately 300 base pairs at the end of *cheB* were kept in the genome to maintain the proper regulation of the downstream expression of *cheY* and *cheZ* (Victor Sourjik, personal communication). The sequences homologous to the targets *(cheR* and *cheB*) in the genome were added to the oligonucleotide primers. PCR reactions were performed with these primers to amplify the sequences containing a tetracycline resistant cassette flanked by flippase recognition target (FRT) sites from pCP16 [70]. Cells were first transformed with the plasmid pKD46, and then transformed with the purified PCR product after induction of the recombinase protein from pKD46. After successful recombination, the portion of the genome containing the deletion and the tetracycline cassette was transduced to a new *E. coli* RP437 background using the phage P1vir. Finally, the tetracycline resistant cassette was excised from the genome with flippase (Flp) by transforming the mutant strain with pCP20 leaving a single FRT sequence scar to obtain the strain NWF121 (Δ*cheRcheB*).

Gene fusions of *cheR* and *cheB* with the genes encoding for the fluorescent reporters sfYFP, [71], or mCherry [72], which have been codon-optimized for *E. coli* expression, and a cassette containing a kanamycin marker flanked by FRT sites (from pCP15 [70]) were constructed using the Gibson assembly method [73] from PCR fragments and a pUC19 vector backbone. The constructs were PCR amplified with sequences homologous to the targets added to the oligo primers and recombined separately into the wild-type MG1655 strain following the same protocol described above. Constructs were recombined into either the native lactose *(lac)* or rhamnose (*rha*) operon loci in the chromosome to take advantage of the host inducible transcription regulation. Each construct was transduced sequentially into the mutant RP437 strain lacking *cheR* and *cheB* (NWF121) using the phage P1vir and excision of the kanamycin resistant cassette with the flippase after each successful transduction. A gene coding for the fluorescent protein sfCFP under the control of the constitutive promoter *pBla* was also recombined into the genome of the mutant strains to provide an independent fluorescence control. Two strains, which are almost identical except that the inducible promoters are swapped, were obtained: YSD2072 (Δ*cheRcheB-FRT, pRha-mCherry-cheR-FRT, pLac-cheB-mYFP-FRT, pBla-CFP-FRT*) and YSD2073 (Δ*cheRcheB-FRT, pLac-mCherry-cheR-FRT, pRha-cheB-mYFP-FRT, pBla-CFP-FRT*). An identical approach was used to clone *pLac-cheB-mYFP* and *pBla-CFP* into RP4972 (Δ*cheB*) to create YSD2044 (Δ*cheB, pLac-cheB-mYFP-FRT, pBla-CFP-FRT*). Deletions and insertions in the final mutant strains were verified by PCR and DNA sequencing.

### Single-cell fluorescence microscopy

Fluorescence microscopy images were acquired using an inverted microscope fitted with a 100x oil immersion objective (Nikon CFI Plan Fluor, N.A. 1.30, W.D 0.2 mm), a solid state white light source (SOLA II SE, Lumencor), a digital scientific CMOS camera (Hamamatsu ORCA-Flash4.0 V2, 1x1 pixel binning, rolling shutter, full frame, 16 bits). Cells were spotted on agarose pads (1% wt/vol agarose with M9 salts) after being washed twice in M9 salts and mounted between a glass slide and a #1.5 glass coverslips. Five different frames containing on average 200 cells were acquired in phase contrast and three fluorescence channels for each sample (CFP filters ex436/20, 455LP, em480/40, YFP filters ex500/20, 515LP, em535/30, mCherry filters ex560/40, 585LP, em630/75). The camera dark current was subtracted from each images and the uneven illumination was corrected using a flat-field image acquired using uniform fluorescent slides.

Cell outlines were determined using MicrobeTracker [48] on the phase contrast images. Cells with sizes deviating from the population by more than three standard deviations were discarded from the analysis. Single-cell fluorescence intensities were calculated by summing the fluorescence signal over each cell area and subtracting the background fluorescence intensity.

The autofluorescence of wild-type cells (RP437) in each channel was determined and subtracted from the fluorescence intensities of cell expressing the fluorescent reporters. The small amount of cross talk between the fluorescent proteins was determined using cells expressing single fluorescent labels and corrected in cells expressing multiple labels using linear unmixing [74].

### Quantitative immunoblotting and fluorescence calibration

The calibration of cell culture optical density (OD_600_) to colony forming units (CFU) was done using serial dilution and plating. Cells expressing different concentrations of the fluorescently labeled proteins were suspended in Laemmli buffer, then boiled for 5 minutes and homogenized in an ultrasonic water bath for 1 minute. Known concentrations of purified fluorescent protein standards (GFP: Rockland 000-001-215 lot 23193, and RFP: abCam ab51993 lot GR25411-12) were added to wild-type cell lysate and treated with the same conditions as the samples to generate standard curves. The lysate of 10^8^ cells in 20 μL were loaded in each lane of pre-casted polyacrylamide gels (BioRad cat. #456-9035) and run in Tris/glycine/SDS buffer at 100 Volts for 90 minutes at 4°C. The proteins were transferred to a low fluorescence 0.45 μm PVDF membrane (BioRad, cat. #162-0261) using wet transfer in Tris/glycine/20% Methanol buffer at 100V for 60 minutes at 4°C. The membranes were blocked to prevent non-specific antibody binding using blocking buffer (EMD Millipore, cat. # WBAVDFL01) for 60 minutes at room temperature. To detect CheB-mYFP, mCFP, and standard GFP, the membranes were hybridized with 1:5,000 dilutions of anti-GFP antibodies conjugated to DyLight488 (Rockland cat. #600-141-215 lot 23518) in Tris buffer saline with 0.05% Tween 20 pH 7.5 for 12 hours with gentle agitation at 4°C. To detect mCherry-CheR and standard RFP, the membranes were hybridized first with 1:2,500 dilutions of anti-RFP antibodies (abCam cat. # ab183628 lot GR170176-1) in Tris buffer saline with 0.05% Tween 20 pH 7.5 for 12 hours with gentle agitation at 4°C, then with 1:10,000 dilutions of anti-Rabbit antibodies conjugated with Dylight488 (Rockland #611-141-002 lot 23521) for 1 hour at room temperature. The membranes were washed three times for 15 minutes with Tris buffer saline with 0.05% Tween 20 pH 7.5 after each incubation. The membranes were dried and scanned with a laser scanner (GE Typhoon 9400) at 488 nm. The images were processed with ImageJ [75] to quantify the signal intensities.

The calibration of fluorescence intensities to protein numbers was calculated using Bayesian linear regression of the quantitative immunoblotting data with the average cell fluorescence signals to obtain the posterior probability distributions of the fluorescence signal per protein. The regression model was setup in the R statistical computing environment [76] with the RStan package [77]. The number of each fluorescent protein per cell was determined as the maximum a posteriori estimate.

### *In situ* hydrogel polymerization and automated cell imaging

Trapping cells with fast *in situ* hydrogel polymerization was done by supplementing the motility buffer with 5% wt/vol polyethylene glycol diacrylate (PEGDA) (M.W ~2,000, JenKem Technology cat. #A4047-5) and 0.05% wt/vol of the photoiniator lithium phenyl-2,4,6-trimethylbenzoylphosphinate [51]. To remove traces of reactive contaminants in the PEGDA, a 20% wt/vol solution was incubated for 10 minutes with a high concentration of washed *E. coli* cells. Cells were removed by centrifugation and filtration through a 0.22 μm filter. The hydrogel polymerization was triggered by exposing the sample for 5 seconds with violet light using a solid state light source (SOLA II SE, Lumencor) at full intensity through a band pass excitation filter (395/25) and the microscope 10X objective (Nikon CFI Plan Fluor, N.A. 0.30, W.D 16.0mm).

The automated imaging of immobilized cells was done using custom Matlab scripts controlling the microscope and a motorized stage (Prior Scientific, cat. # H117) through the Micro-Manager core library [78] (the code is available at https://github.com/dufourya/FAST). After immobilization, the cell coordinates were registered using image analysis in Matlab and the microscope was configured for epifluorescence imaging at 100X. The computer-controlled stage moved sequentially to each cell location. The z-focus was automatically adjusted for each cell before imaging in phase contrast and the three fluorescence channels (CFP filters ex436/20, 455LP, em480/40, YFP filters ex500/20, 515LP, em535/30, mCherry filters ex560/40, 585LP, em630/75). Cells that were not properly aligned with the focal plane, which were determined by detecting non-closed edges of the outlines of cells in the analysis of phase contrast images, were skipped. Cells with sizes deviating from the population by more than three standard deviations were discarded from the analysis. About 200 cells were imaged in less than 40 minutes for each experiment trial. The fluorescence signal from each cell did not change significantly as a function of time during the single-cell imaging phase indicating that the fluorescent protein fusions are stable when the cells are trapped in the hydrogel (S8 Fig.).

### Modeling individual cell tumble bias as a function of chemotaxis protein numbers

The model used to calculate tumble bias as a function of protein numbers is based on a previously published model [12]. The concentration of phosphorylated CheA [*CheA_P_*] changes according to

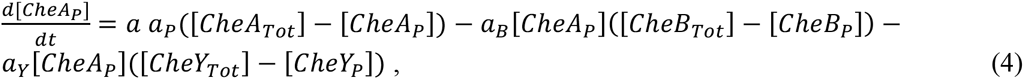

inwhich *a* is the receptor cluster activity, [*CheA_Tot_*], [*CheB_Tot_*], [*CheY_Tot_*], and [*CheY_P_*] are the concentrations of all CheA, all CheB, all CheY, phosphorylated CheY, *a_P_, a_B_*, and *a_Y_* are the phosphorylation rate constants.

The concentrations of phosphorylated CheB and CheY changes according to

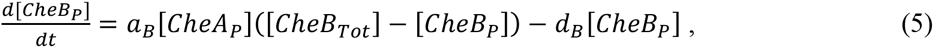

and

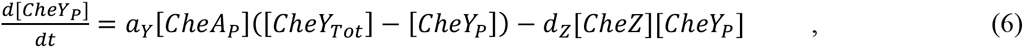

in which *d_B_* and *d_Z_* are the dephosphorylation rates and *[CheZ]* is the concentration of CheZ. The biochemical rate and binding parameters are kept the same for all cells (the values used are summarized in S4 Table). To simulate cell-to-cell variability in protein expression, the protein numbers were sampled for each cell from log-normal distributions according to a noisy gene expression model as previously described [12] using experimentally determined average concentrations [52]. To match as well as possible our experimental results, the intrinsic and extrinsic noise levels in the expression of all the chemotaxis proteins (except CheR and CheB which were measured directly) were reduced to be a tenth of what was previously proposed [12]. The concentration of CheZ was reduced slightly to match the observed range of tumble bias in our experiments. In the absence of stimuli, equation 1–6 can be solved at equilibrium to calculate the steady-state concentration of each protein.

The variance of [*CheY_P_*] resulting from the spontaneous fluctuations of the cluster activity for each single cell was calculated using the linear noise approximation of the Master equation as previously described for the chemotaxis system [45]. Briefly, taking the stoichiometry matrix of the system, *S*, and the propensity vector, **ν**, the diffusion matrix of the system, *B*, was calculated using the linear noise approximation by solving *B^T^B = S* diag(**ν**)S^T^ [79, 80]. The correlation matrix, *C*, which contains the variance and covariance for the fluctuations of all the components in the model, was calculated from the linearized rate equations near the equilibrium solution given by the Jacobian matrix of the system, *A*, by solving numerically the Lyapunov equation *AC* + *CA^T^* + *B^T^B* = 0 [81, 82].

To simulate the effect of the slow fluctuations in cluster activity on the concentration of phosphorylated CheY, [*CheY_P_*] was sampled randomly from a Gaussian distribution centered at the steady-state [*CheY_P_*] with variance var([*CheY_P_*]) form the correlation matrix, *C*. Because the cluster activity fluctuates according to the adaptation time scale, the effective [*CheY_P_*] was calculated from the average of *n_eff_* samples according to 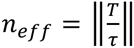, in which T is the average length of the recorded trajectory in seconds (100 seconds) and τ is the slowest relaxation time-scale of the Jacobian matrix *A* evaluated at the equilibrium solution given by the largest of the eigenvalues λ of *A, τ* = —1/max (λ). The tumble bias was calculated as a function of the effective [*CheY_P_*] using the steady-state function of the adaptive flagellar motor as previously described [13] and the coordination of multiple flagella as previously described [34].

## Acknowledgements

We thank Adam Waite and Setsu Kato for comments on the manuscript. We thank Victor Sourjik and Sandy Parkinson for their generous gifts of strains. This work was supported by the Paul Allen foundation (award no. 11562), and by the National Institute of Health (R01GM106189).

## Supporting Information

**S1 Table.** Table of plasmids used in this study.

| Name | Origin of replication | Marker | Promoter | Gene(s) of interest | Description | Reference |
| --- | --- | --- | --- | --- | --- | --- |
| pKD46 | repA101ts | AmpR | pAraB | *gam, bet, exo* | Temperature sensitive plasmid containing the Phage $\lambda$ Red recombination system | (Datsenko & Wanner, 2000) |
| pCP15 | pMB1 | AmpR, Kan R |  | *FRT-KanR-FRT* | Kanamycin resistance cassette flanked by FRT sequences | (Cherepanov & Wackernagel, 1995) |
| pCP16 | pMB1 | AmpR, Tet R |  | *FRT-tetAR-FRT* | TetRacycline resistance cassette flanked by FRT sequences | (Cherepanov & Wackernagel, 1995) |
| pCP20 | repA101ts | AmpR, Cm R |  | *flp* | Temperature sensitive plasmid expressing the FLP recombinase ("flippase") | (Cherepanov & Wackernagel, 1995) |
| pTU136 | R6K | AmpR | pLac | *ssdsbA-mCherry* | Template for mCherry gene sequence | (Uehara, Dinh, & Bernhardt, 2009) |
| pYSD1003 | pMB1 | AmpR |  | *sfYFP* | Template for super-folder mYFP gene sequence | This study |
| pYSD1004 | pMB1 | AmpR |  | *sfCFP* | Template for super-folder mCFP gene sequence | This study |
| pYSD1011 | pMB1 | KanR | pBla | *sfCFP* | mCFP under the control of the *bla* promoter from pUC19 | This study |
| pYSD1007 | pMB1 | AmpR, Kan R | pLac | *cheB-mYFP* | Template for the translational fusion of CheB and mYFP | This study |
| pYSD1005 | pMB1 | AmpR, Kan R | pLac | *mCherry-cheR* | Template for the translational fusion of mCherry and CheR | This study |

**S2 Table.** Table of bacterial strains used in this study.

| Name | Description | Reference |
| --- | --- | --- |
| RP437 | Wild-type for chemotaxis | (Parkinson, 1978) |
| RP4972 | ΔcheB | (Parkinson, 1978) |
| NWF121 | ΔcheRcheB-FRT with ~300 bp CheB 3' fragment | This study |
| YSD2023 | pLac-mCherry-cheR, FRT-kanR-FRT | This study |
| YSD2024 | pRha-mCherry-cheR, FRT-kanR-FRT | This study |
| YSD2025 | pLac-cheB-mYFP, FRT-kanR-FRT | This study |
| YSD2027 | pRha-cheB-mYFP, FRT-kanR-FRT | This study |
| YSD2031 | pBla-mCFP, FRT-kanR-FRT | This study |
| YSD2040 | ΔcheRcheB, FRT, pLac-cheB-mYFP, FRT | This study |
| YSD2041 | ΔcheRcheB, FRT, pLac-mCherry-cheR, FRT | This study |
| YSD2044 | ΔcheB, pLac-cheB-mYFP, FRT | This study |
| YSD2062 | ΔcheRcheB, FRT, pLac-cheB-mYFP, FRT, pRha-mCherry-cheR, FRT | This study |
| YSD2063 | ΔcheRcheB, FRT, pRha-cheB-mYFP, FRT, pLac-mCherry-cheR, FRT | This study |
| YSD2072 | ΔcheRcheB, FRT, pLac-cheB-mYFP, FRT, pRha-mCherry-cheR, FRT, pBla-mCFP, FRT | This study |
| YSD2073 | ΔcheRcheB, FRT, pRha-cheB-mYFP, FRT, pLac-mCherry-cheR, FRT, pBla-mCFP, FRT | This study |

**S3 Table.**
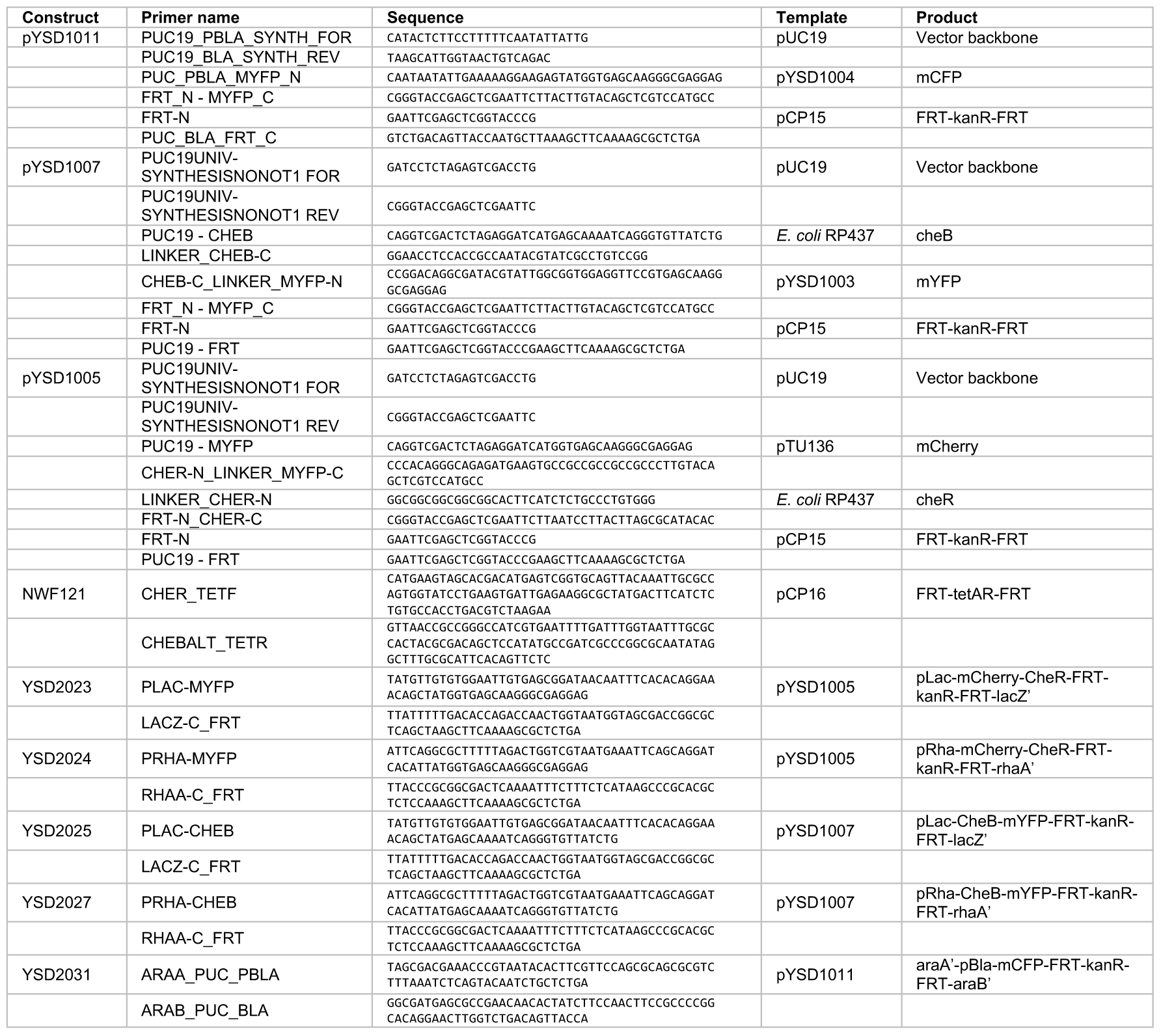
Table of oligonucleotides primers used in this study.

**S4 Table.**
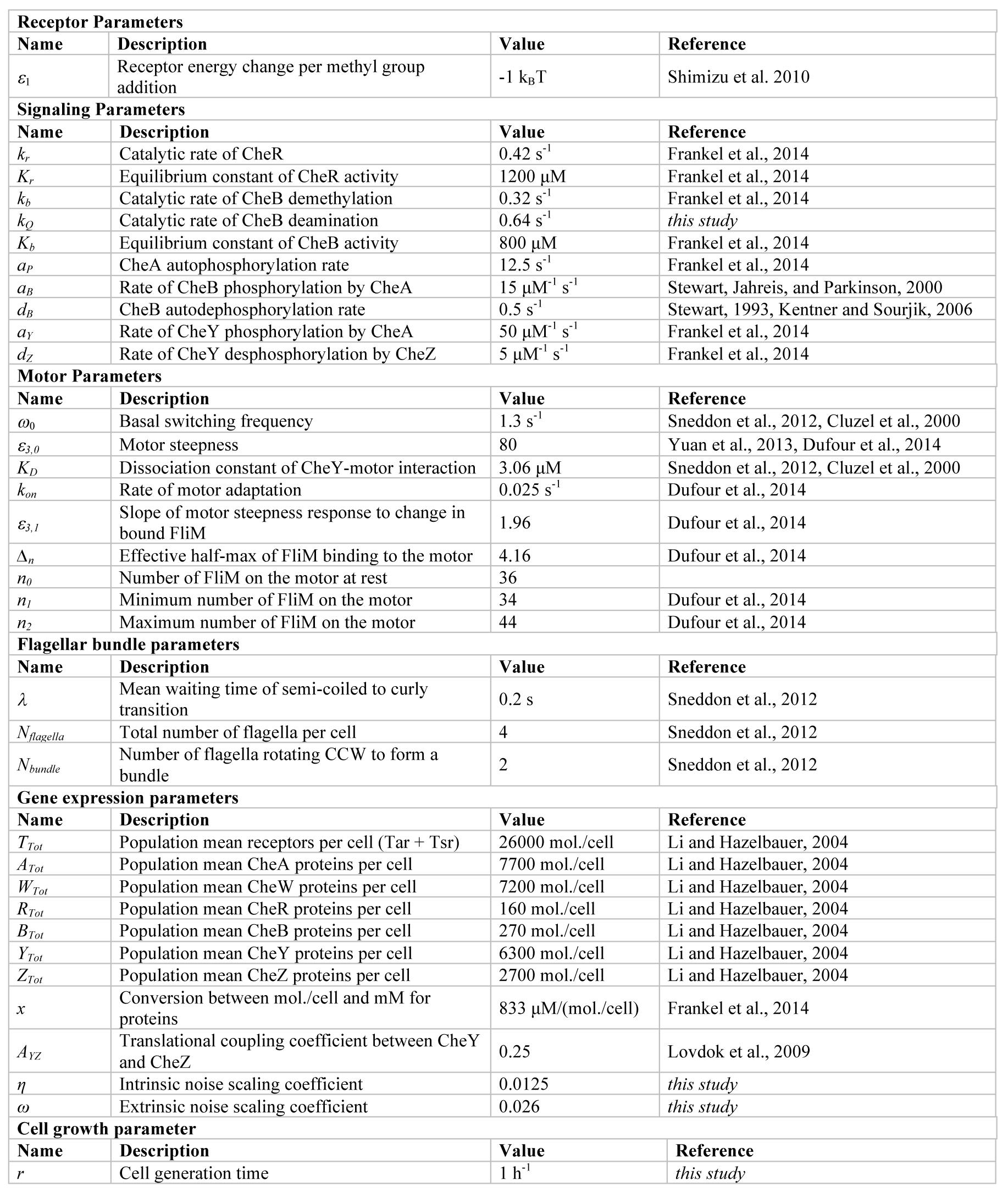
Table of model parameters used in this study.

**S1 Fig.**
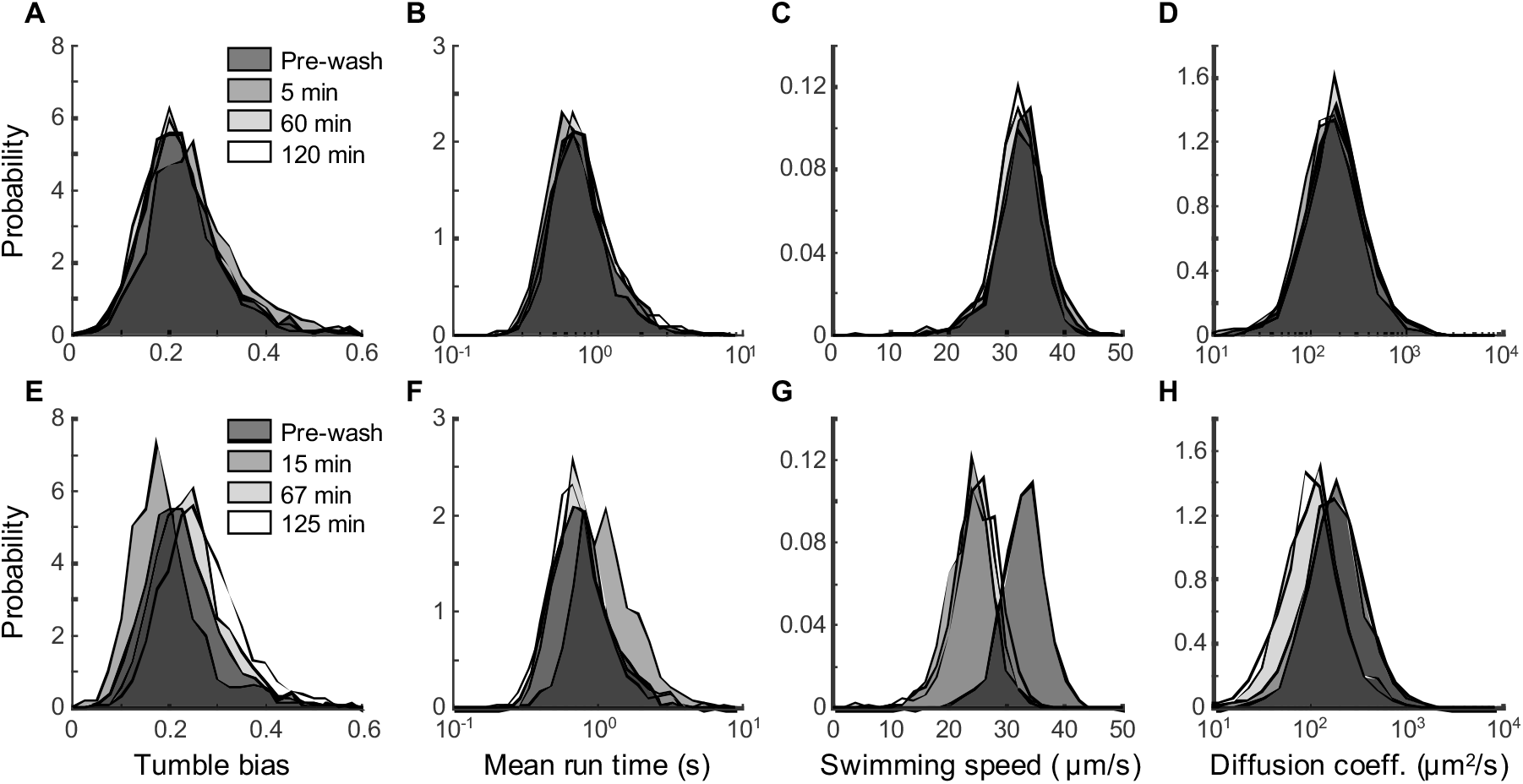
Effect of the chemotaxis buffer, PEGDA, and LAP on the distributions of swimming phenotypes in a clonal *E. coli* RP437 population. (A-D) Probability distribution of cell tumble biases, mean run times, mean swimming speeds, and cell diffusion coefficients from cells swimming in chemotaxis buffer after the indicated incubation times. (E-H) Same distributions from cells swimming in chemotaxis buffer supplemented with 5% PEGDA and 0.05%LAP after the indicated incubation times.

**S2 Fig.**
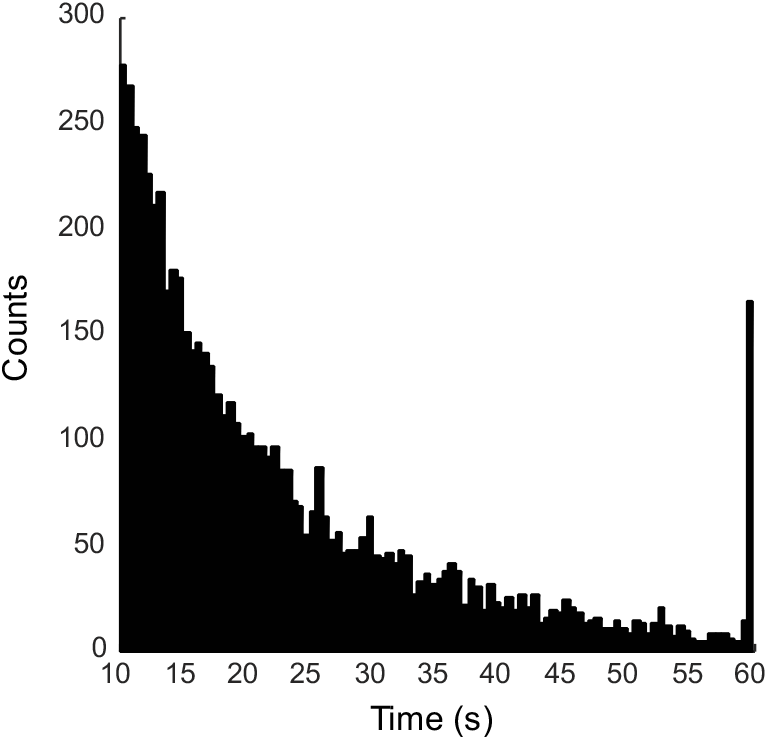
Distribution of trajectory length obtained from tracking 6,332 individual swimming RP437 cells for 60 seconds.

**S3 Fig.**
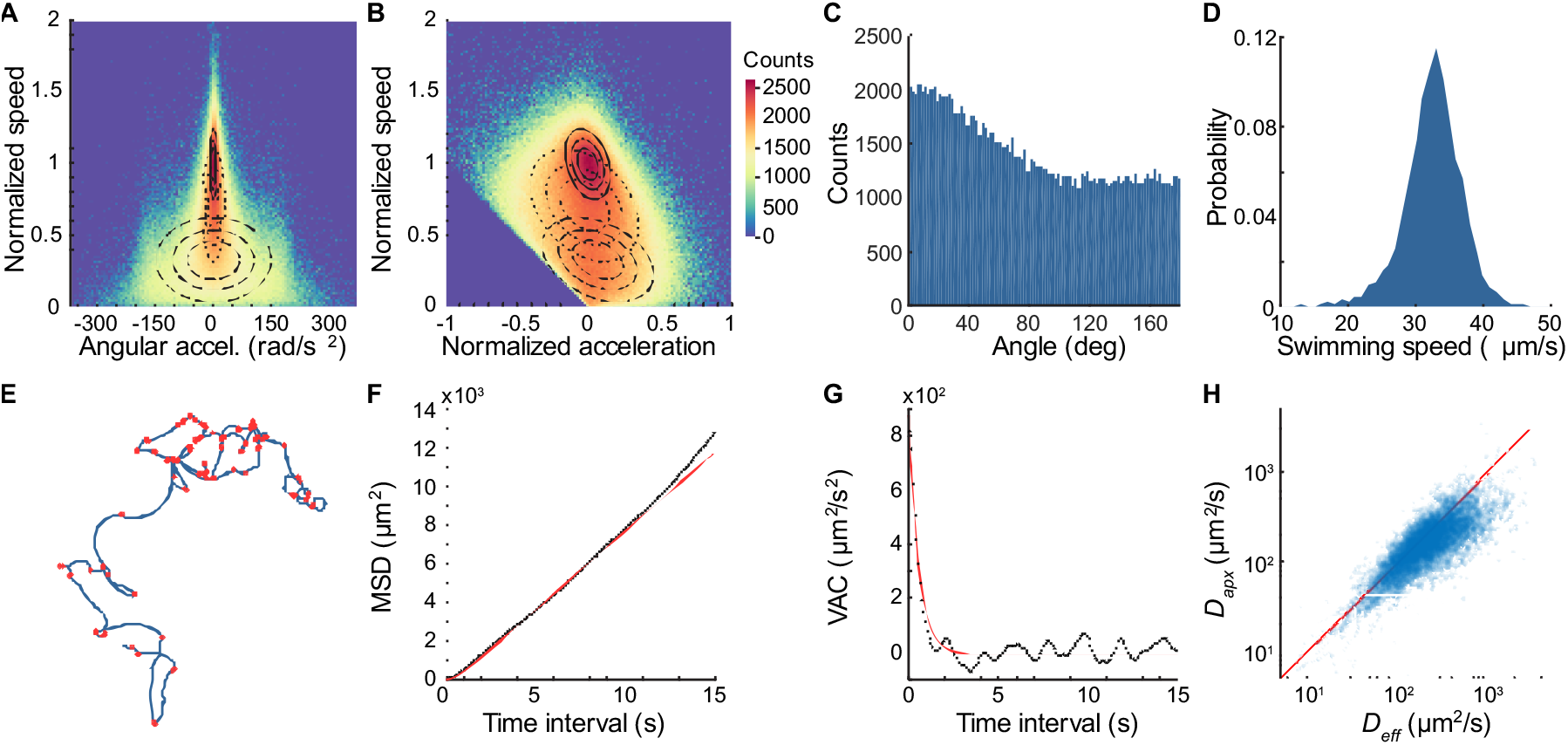
Tumble detection and diffusion coefficient calculations. (A) Density plot of normalized cell swimming speed as a function of angular acceleration. (B) Density plot of normalized cell swimming speed as a function of normalized cell acceleration. The three-dimensional density distribution comprising ~6 millions data points was fitted with a mixture of three tri-variate Gaussian distributions to represent three possible cell swimming states: running (solid lines), tumbling (dashed lines), and intermediate (dotted lines). (C) Distribution of angles measured from the change in direction in the swimming trajectories after each detected tumble for RP437 cells. (D) Probability distribution the mean swimming speeds of individual cells. (E) Example of a 60 seconds single-cell trajectory where detected tumbles are marked with red dots. (F) Mean square displacement and (G) velocity auto-correlation as a function of time intervals calculated from a representative cell trajectory (black) with the corresponding fit (red) to extract the cell diffusion coefficient. (H) Scatter plot of the approximated diffusion coefficients (*D_apx_*) calculated from the mean run time between tumbles against the effective diffusion coefficient (*D_eff_*) calculated for the cell directional persistence for each cell. The distributions were calculated from about 6,000 individual trajectories combined from three independent experiments.

**S4 Fig.**
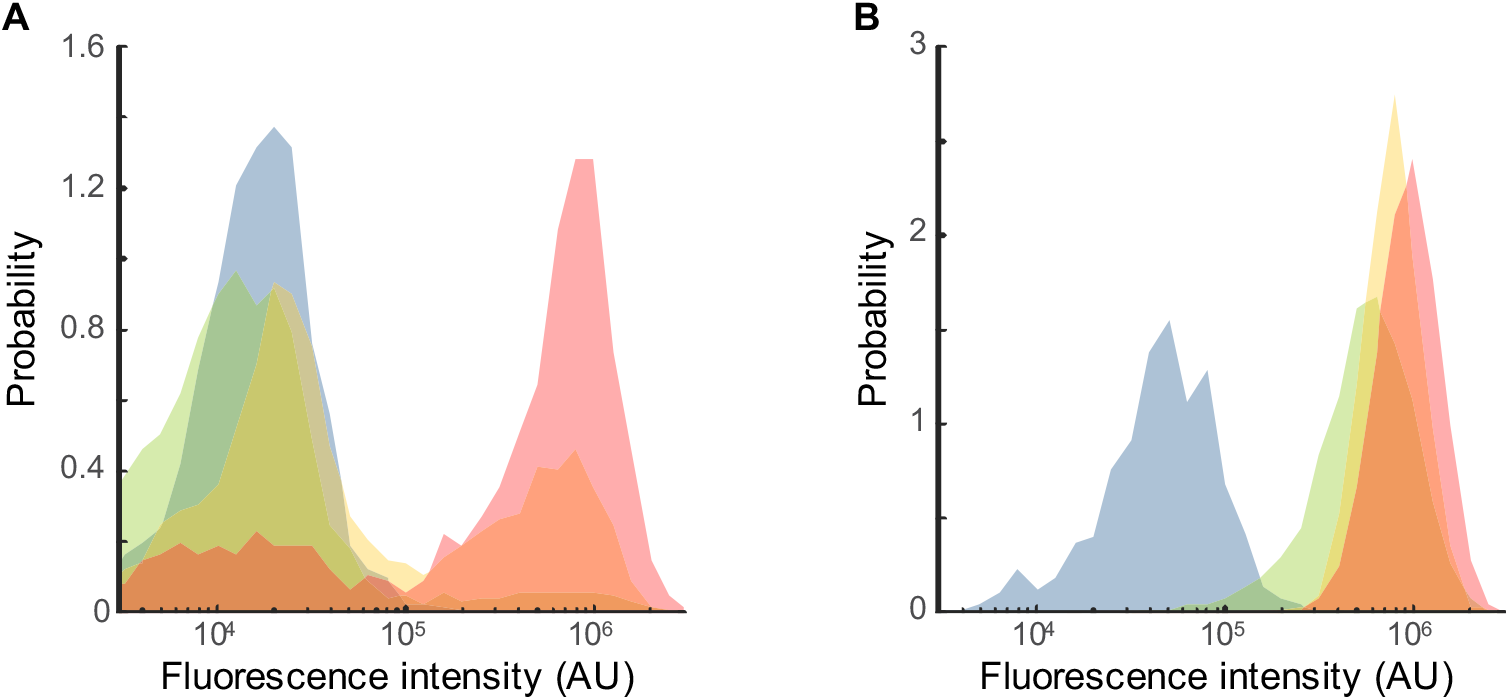
Induction of fluorescently labeled chemotaxis proteins. (A) Probability distributions of fluorescence intensities from the inductions of CheB-mYFP in the YSD2073 mutant strain (pRha cheB-mYFP, pLac mCherry-cheR) with four rhamnose concentrations: 0 mM (cyan), 0.3 mM (green), 1 mM (yellow), and 3 mM (red). (B) Probability distributions of fluorescence intensities from the inductions of mCherry-CheR in the same strain with four IPTG concentrations: 0 μM (cyan), 10 μM (green), 30 μM (yellow), and 100 μM (red). The fluorescence intensities were obtained from the analysis of thousands of cells using MicrobeTracker on epi-fluorescence microscopy images. The bimodal distribution of fluorescence intensities from the expression of CheB-mYFP is a result of the bi-stability of the pRha promoter.

**S5 Fig.**
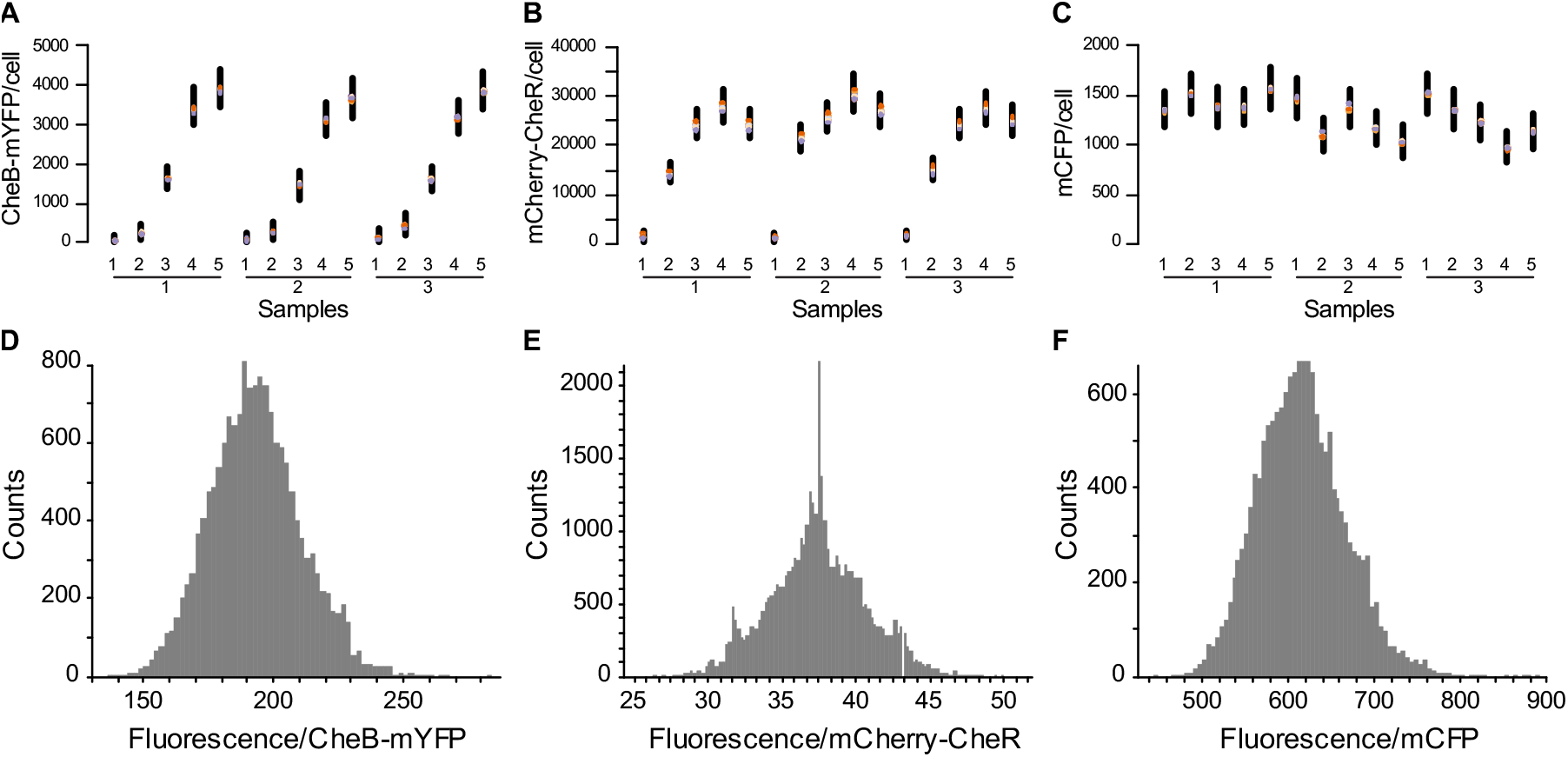
Calibration of fluorescence units per fluorescent molecules using Bayesian regression analysis with the YSD2073 mutant strain (pRha cheB-mYFP, pLac mCherry-cheR, pBla mCFP). (A) Estimated numbers of CheB-mYFP molecules per cell as a result of induction with different rhamnose concentrations (1: 0 mM, 2: 0.3 mM, 3: 1 mM; 4: 3 mM, 5: 10 mM) shown for three independent experiments. The black lines represent the 80% confidence interval. The colored dots indicate the median for each of the 8 chains of the MCMC sampling. (B) Estimated numbers of mCherry-CheR molecules per cell as a result of induction with different IPTG concentrations (1: 0 μM, 2: 10 μM, 3: 30 μM; 4: 100 μM, 5: 1 mM) shown for three independent experiments. (C) Estimated numbers of mCFP molecules per cell as a result of induction with different IPTG concentrations (1: 0 μM, 2: 10 μM, 3: 30 μM; 4: 100 μM, 5: 1 mM) shown for three independent experiments. As expected mCFP expression does not respond to the presence of IPTG or rhamnose. (D) Posterior probability distribution of the expected number of fluorescence units per CheB-mYFP molecule. (E) Posterior probability distribution of the expected number of fluorescence units per mCherry-CheR molecule. (F) Posterior probability distribution of the expected number of fluorescence units per mCFP molecule.

**S6 Fig.**
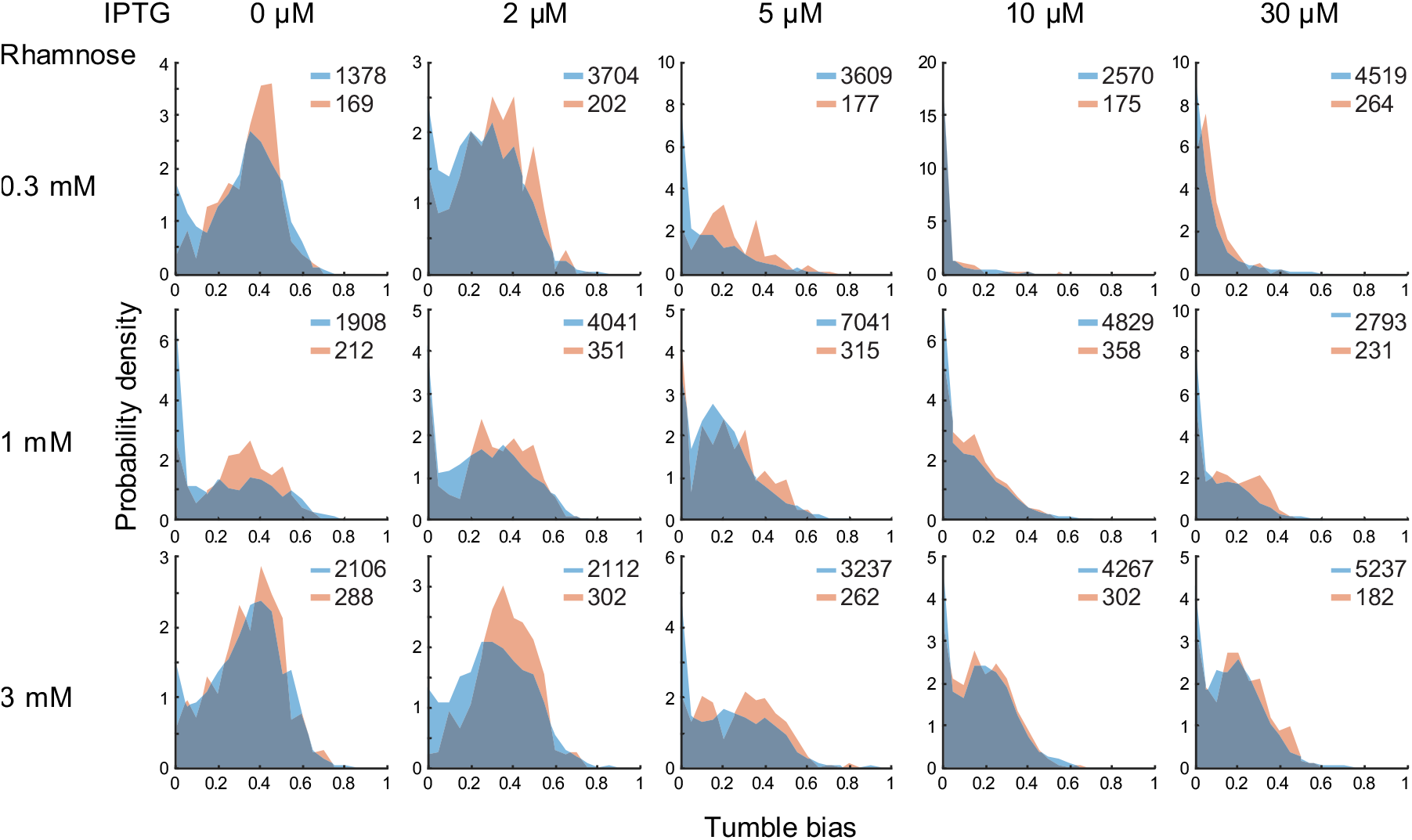
Manipulating and sampling tumble bias distributions in a mutant *E. coli* strain expressing mCherry-CheR and CheB-mYFP. The YSD2072 mutant strain (pLac cheB-mYFP, pRha mCherry-cheR, pBla mCFP) was grown in M9 glycerol medium supplemented with the indicated concentrations of the inducers rhamnose and IPTG to obtain different distributions of tumble biases. The distributions of phenotypes from the population of cells trapped and imaged in the hydrogel (red) is comparable to the distribution of phenotypes from the entire cell population (blue) indicating that the trapped cells represent an unbiased sample of the population. The number of cells represented in each distribution is indicated for each plot.

**S7 Fig.**
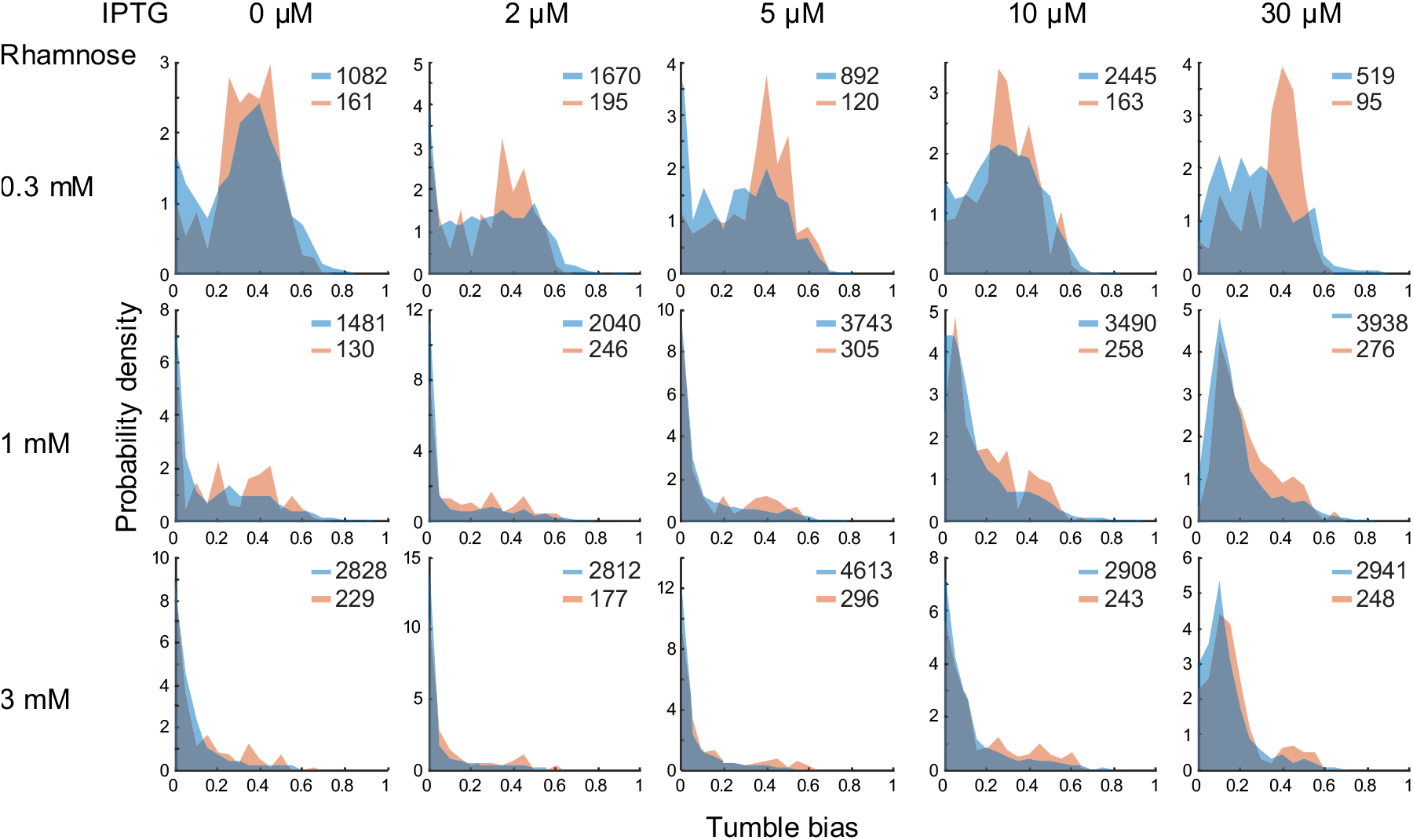
Manipulating and sampling tumble bias distributions in a mutant *E. coli* strain expressing mCherry-CheR and CheB-mYFP. The YSD2073 mutant strain (pRha cheB-mYFP, pLac mCherry-cheR, pBla mCFP) was grown in M9 glycerol medium supplemented with the indicated concentrations of the inducers rhamnose and IPTG to obtain different distributions of tumble biases. The distributions of phenotypes from the population of cells trapped and imaged in the hydrogel (red) is comparable to the distribution of phenotypes from the entire cell population (blue) indicating that the trapped cells represent an unbiased sample of the population. The number of cells represented in each distribution is indicated for each plot.

**S8 Fig.**
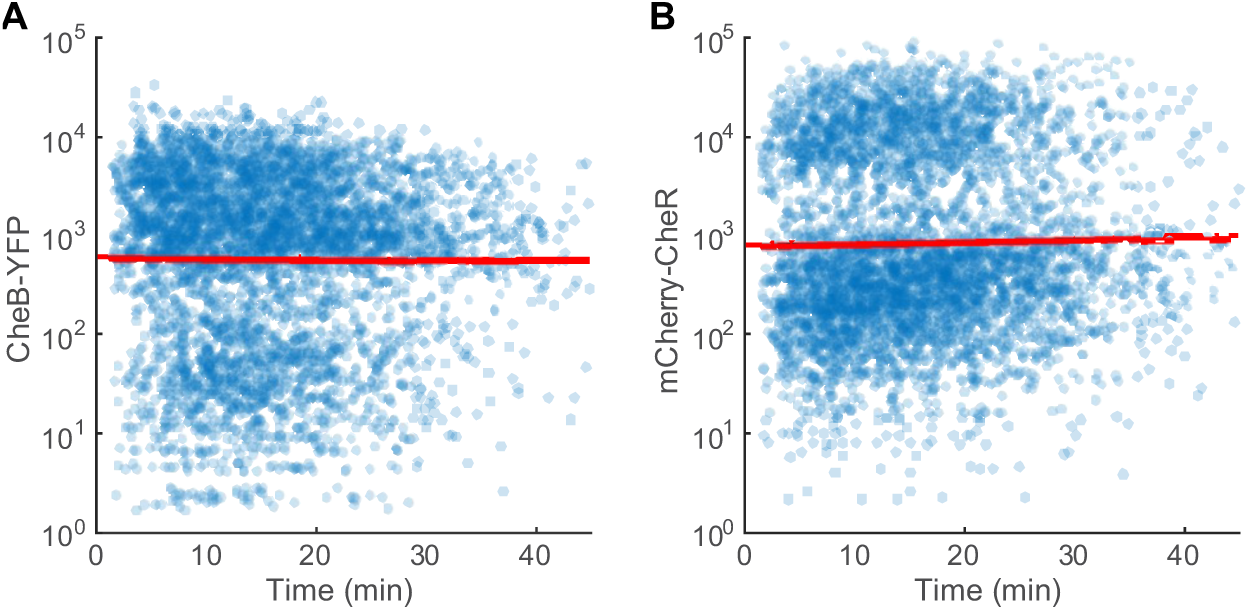
Protein stability during single-cell fluorescence imaging of cells immobilized in the hydrogel. (A) Scatter plot of the estimated number of CheB-YFP proteins in each cell as a function of time after cell immobilization. A linear fit (red line) indicates that there is no significant change in protein numbers as a function of time (slope −0.0022 min^−1^, 95% confidence interval [-0.0094; 0.0050]). (B) Scatter plot of the estimated number of mCherry-CheR proteins in each cell as a function of time after cell immobilization. A linear fit (red line) indicates that there is no significant change in protein numbers as a function of time (slope 0.0049 min^−1^, 95% confidence interval [-0.0025; 0.0123]).

**S9 Fig.**
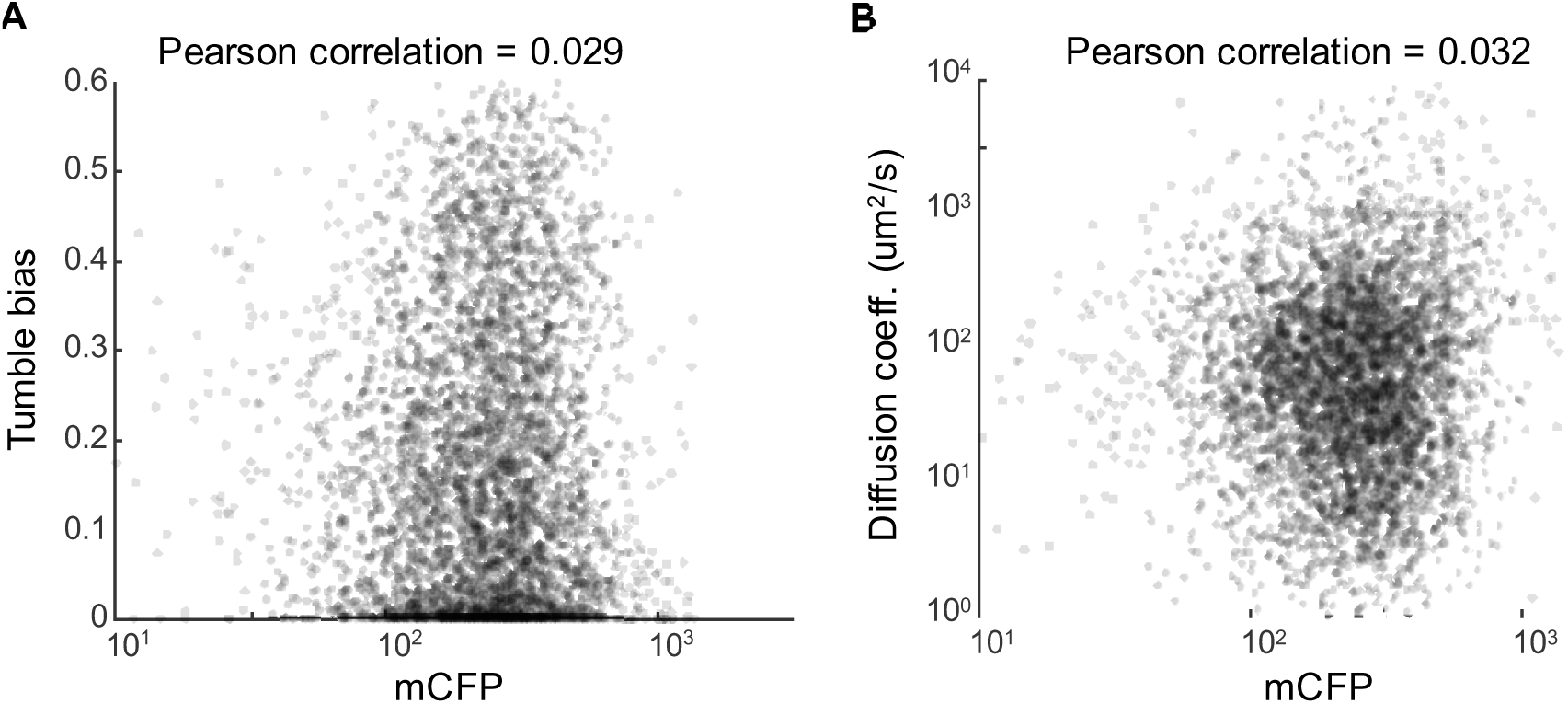
Correlations of single-cell swimming phenotypes with mCFP numbers. (A) Scatter plot of singlecell tumble biases against mCFP numbers. (B) Scatter plot of single-cell diffusion coefficients against mCFP numbers.

**S10 Fig.**
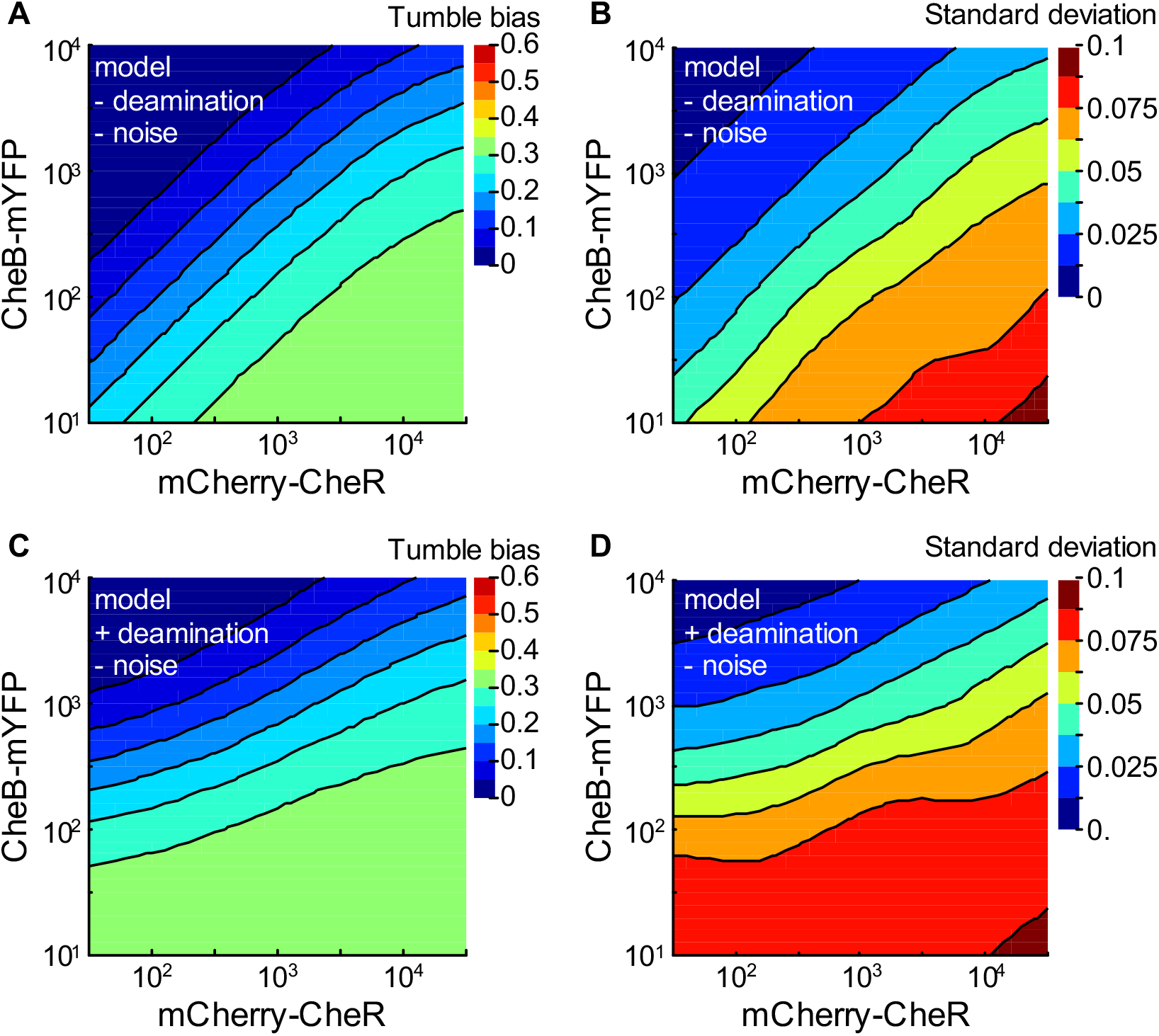
Tumble bias and residual standard deviation as a function of CheR and CheB numbers predicted from a model missing CheB-dependent receptor deamidation and/or receptor adaptation noise. (A) Contour plot of the local linear regression of the predicted tumble bias as a function of CheR and CheB numbers for a model missing both CheB-dependent receptor deamidation and receptor adaptation noise. (B) Contour plot of the predicted residual tumble bias standard deviation resulting from stochastic expression of the chemotaxis proteins with no signaling noise from the receptor cluster. (C) Contour plot of the local linear regression of the predicted tumble bias as a function of CheR and CheB numbers for a model including the deamidation reaction but missing receptor adaptation noise. (D) Contour plot of the predicted residual tumble bias standard deviation resulting from stochastic expression of the chemotaxis proteins with no signaling noise from the receptor cluster. From the stochastic gene expression model, we sampled 8405 cells covering the full range of CheR and CheB expression levels. We then calculated the corresponding tumble bias for each individual cell using a model of bacterial chemotaxis that does not take into account CheB-dependent receptor deamidation or receptor adaptation noise. The local linear regressions were done using a bandwidth of 20% of the data points.

**S11 Fig.**
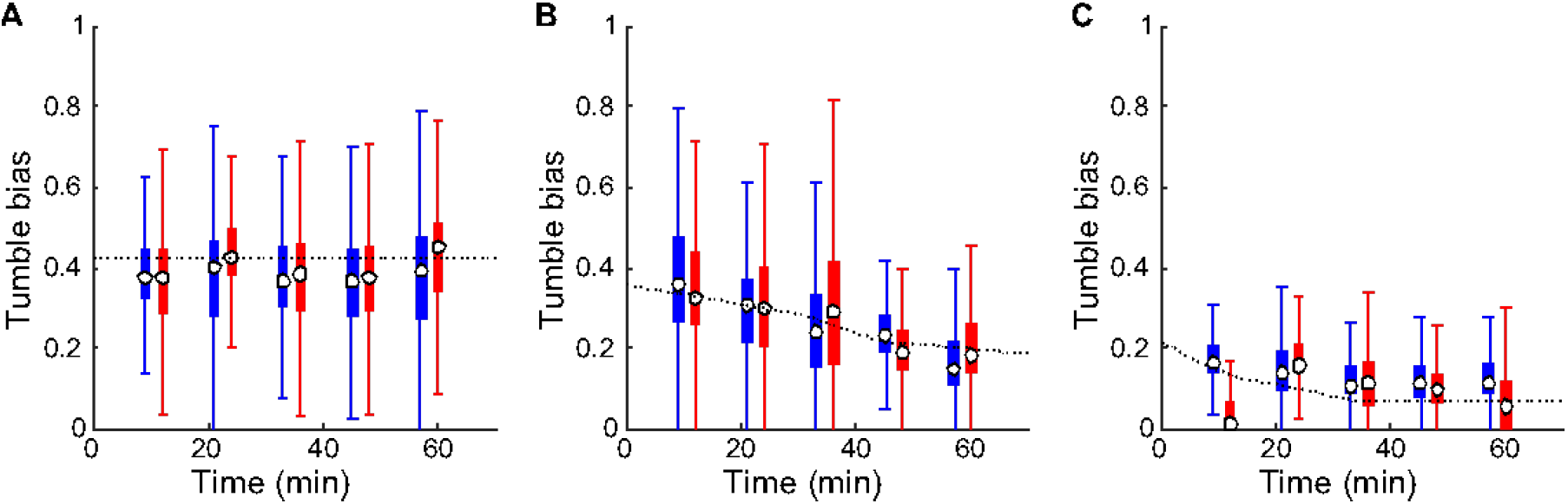
Effect of CheB-YFP expression on the tumble bias in populations of motile cells. Box plots representing the evolution of the distributions of the tumble bias as a function of time of YSD2044 mutant cells grown (A) in the absence of inducer, (B) in the presence of 5 μM IPTG, or (C) in the presence of 50 μM IPTG, after the growth medium was exchanged with chemotaxis buffer. Two independent experimental trials are represented in red and blue. The white dots represent the medians. The boxes span the first and third quantiles. The whiskers indicate 1.5 times the interquartile range. The dotted lines represent the model predictions used to estimate the CheB-dependent deamidation rate (*k_Q_* = 0.64s^−1^) and the number of expressed CheB proteins (0, 100, 250 molecules, respectively). The tumble bias trajectories were calculated over time by first solving the model at steady state with a cell doubling rate set to 1h^−1^ and then numerically solving the evolution of the differential equations after the doubling rate was set to 0 to simulate the transfer of cells from the growth medium to the chemotaxis buffer.

**S12 Fig.**
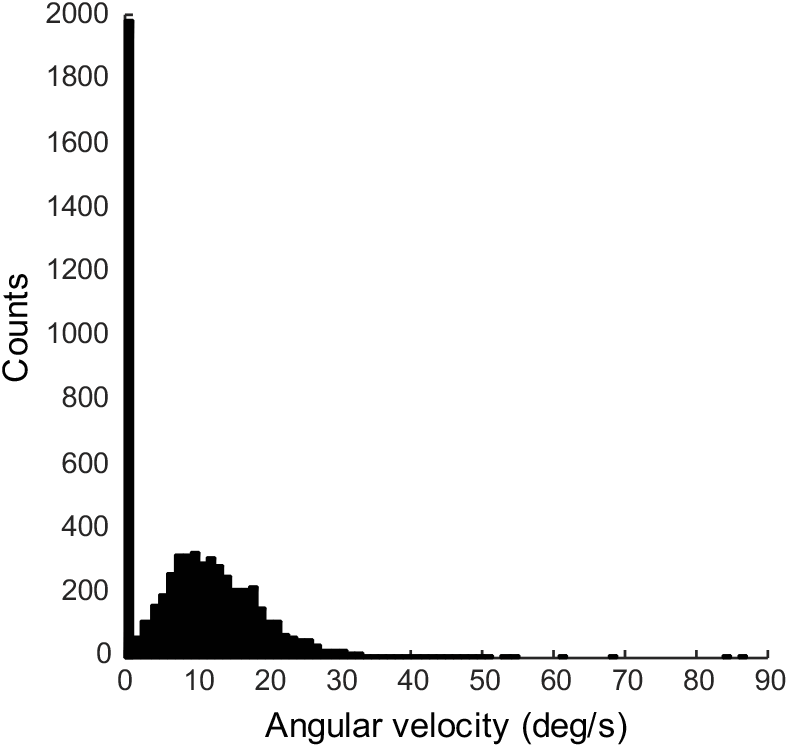
Distribution of angular velocities extracted from the analysis of the mean square displacement and the velocity autocorrelation of each cell trajectory. The angular velocity is non-zero when cells swim close enough to the glass surface to be affected by hydrodynamic interactions between the glass and the cell body rotation.

**S13 Fig.**
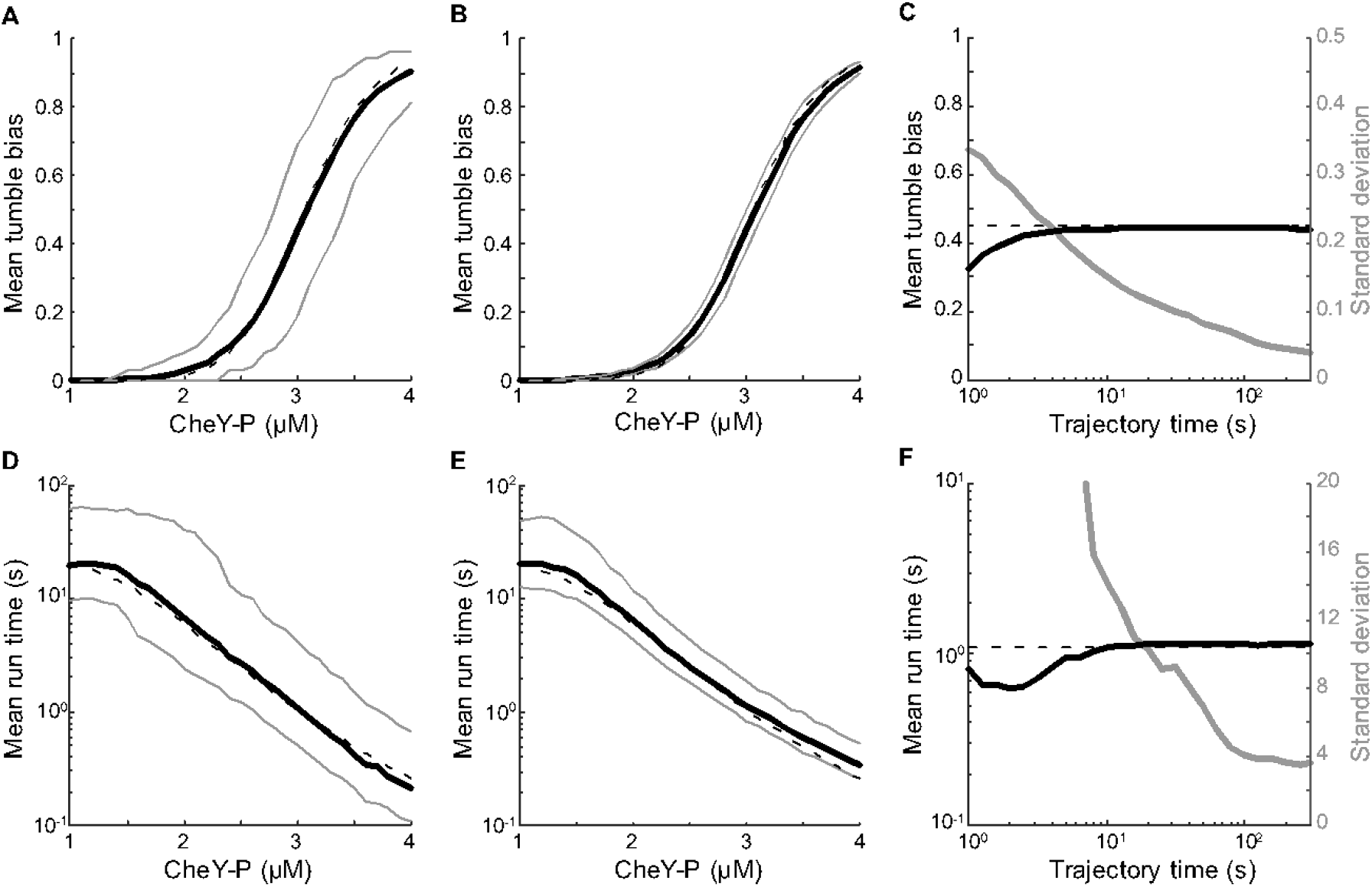
Precision and accuracy of tumble detection on simulated swimming cell trajectories. (A) Mean cell tumble bias as a function of phosphorylated CheY (CheY-P) concentration for 10-second-long trajectories. The dashed line represents the expected theoretical relationship. The tumble detection algorithm was run on 1,000 simulated trajectories for each CheY-P concentrations. The solid line represents the mean of the calculated tumble biases. The grey lines delimit 90% of the probability density. (B) Same as A but for 300 seconds long trajectories. (C) Mean of the calculated tumble bias as a function of total simulation time. The trajectories of cells with 3 μM CheY-P were simulated for different amounts of time between 1 and 300 seconds. The dashed line represents the expected theoretical tumble bias. The tumble detection algorithm was run on 1,000 simulated trajectories for each simulation time. The solid line represents the mean of the calculated tumble biases. The grey line represents the standard deviation of the calculated tumble bias distribution. (D) Mean cell run time as a function of phosphorylated CheY concentration for 10-second-long trajectories. (E) Same as D but for 300 seconds long trajectories. (F) Mean of the calculated mean run time as a function of total simulation time.

